# Neuron-specific protein expression at striatal excitatory synapses

**DOI:** 10.1101/2023.03.16.532957

**Authors:** Yi-Zhi Wang, Tamara Perez-Rosello, D. James Surmeier, Jeffrey N. Savas

**Affiliations:** Department of Neurology, Feinberg School of Medicine, Northwestern University, Chicago, IL 60611, USA; Department of Neuroscience, Feinberg School of Medicine, Northwestern University, Chicago, IL 60611, USA

## Abstract

Combinatorial expression of postsynaptic proteins underlies synapse diversity within and between neuron types ^1-5^. Thus, characterization of neuron-type-specific postsynaptic proteomes is key to obtaining a deeper understanding of discrete synaptic properties and how selective dysfunction manifests in synaptopathies ^6-9^. To overcome the limitations associated with bulk measures of synaptic protein abundance ^6,10^, we developed a biotin proximity protein tagging probe to characterize neuron-type-specific postsynaptic proteomes *in vivo*. We found Shank3 protein isoforms are differentially expressed by direct and indirect pathway spiny projection neurons (dSPNs and iSPNs). Studies in mice lacking Shank3 gene exons 13-16 revealed a robust postsynaptic proteome alteration in iSPNs. We report unexpected cell-type specific synaptic protein isoform expression which could play a key causal role in specifying synapse diversity and selective synapse dysfunction in synaptopathies.

## Main

Postsynaptic specializations are essential for receiving and processing synaptic signals. The composition of postsynaptic proteomes varies from one brain region to the next, reflecting their functional diversity ^1-4^. This diversity is likely to extend to different cell types within brain regions. The striatum is a large subcortical structure involved in goal-directed actions and habits ^11,12^. The principal neurons of the striatum are GABAergic spiny projection neurons (SPNs).

SPNs can be divided into direct and indirect pathway SPNs (dSPNs and iSPNs, respectively) on the basis of their axonal projections, expression of G-protein coupled receptors, and expression of releasable peptides ^11-13^. The extra-striatal innervation of SPNs in the dorsal striatum is derived primarily from glutamatergic neurons in the cerebral cortex and thalamus, which form axospinous synapses. In addition, SPNs are innervated by other SPNs and by GABAergic and cholinergic interneurons ^11-13^.

Striatal dysfunction plays a major role in numerous neurodevelopmental and degenerative diseases such as Huntington’s disease (HD) ^14^, Parkinson’s disease (PD) ^15^ and autism spectrum disorder (ASD) ^16^. Moreover, in each of these disorders, SPN-specific alterations in synaptic function play an important role in the emergence of symptoms ^7-9^. However, the molecular basis of these cell-type specific synaptic changes remains largely unknown since traditional bulk protein methods lack cellular resolution ^6^.

To quantitatively profile cell-specific postsynaptic nano-environments, we designed a BirA* (postBirA*) probe to tag postsynaptic proteins with biotin *in vivo* **(Fig. 1a & Extended Data Fig. 1a)**. We also constructed two negative control BirA* probes that localize to the presynaptic membranes or the cytosol, preBirA* and cytoBirA*, respectively **(Extended Data Fig. 1a)**. We then sub-cloned all three BirA* probes into FLEX plasmids, allowing cell-type specific expression using adeno-associated virus (AAVs) vectors in transgenic mice expressing Cre recombinase under the control of cell-type specific promoters **(Extended Data Fig. 1b)**.

**Fig. 1.**
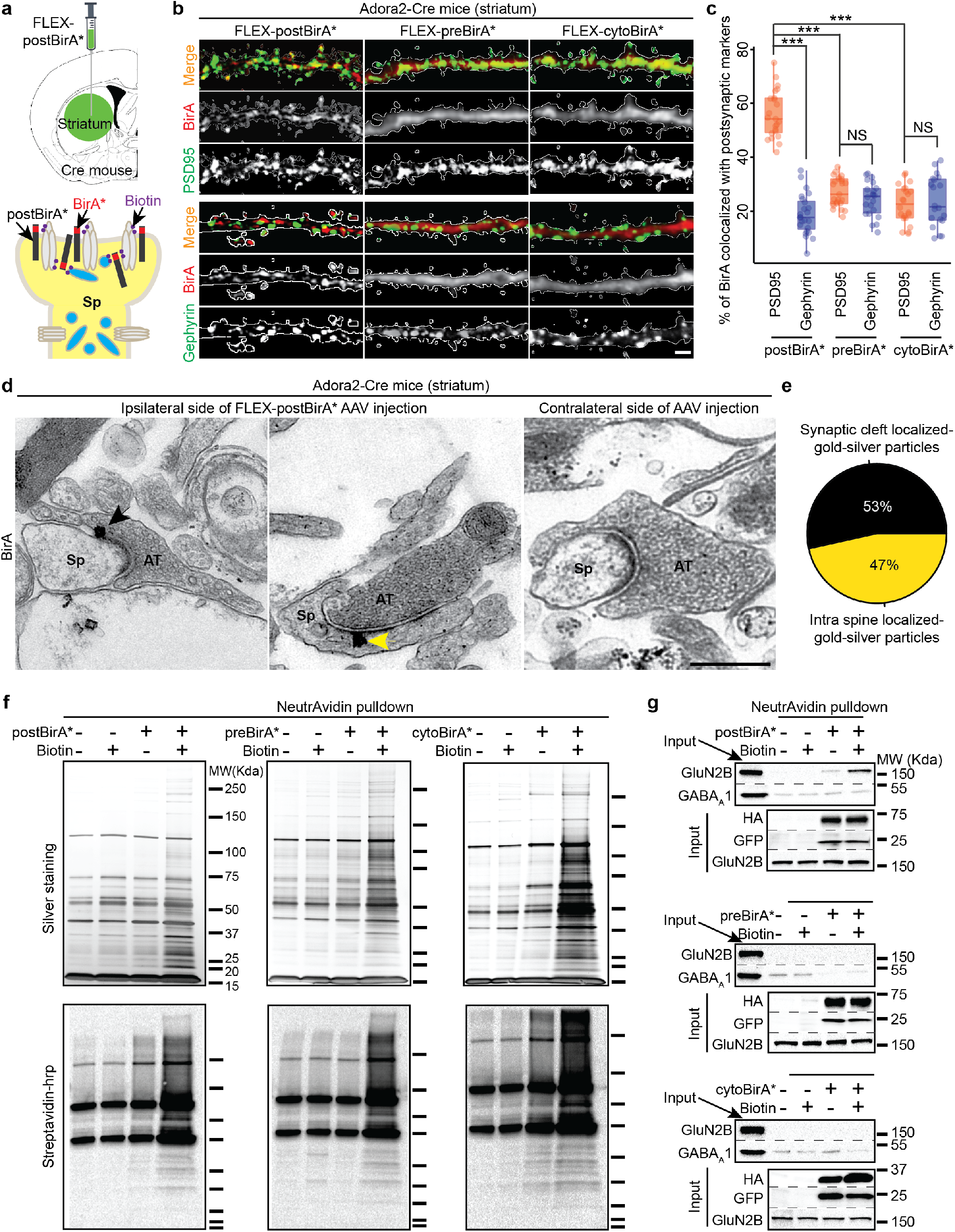
Assessments of postBirA*-based *in vivo* proximity biotin-tagging toolkit. **(a)** Schematic showing intracranial injection of FLEX-postBirA* AAVs into striatum of Cre-mouse (top). Cartoon schematic of postsynaptic localization and proximity biotinylation of postBirA* (bottom). **(b)** Representative IHC analysis showing postBirA*, preBirA*, and cytoBirA* localization relative to the postsynaptic markers PSD95 or Gephyrin. BirA* expressing SPNs were identified based on co-expression of eGFP (white traces). Scale bar, 2 μm. **(c)** Quantification of **(b)**. n = 4-6 mice, 3-5 brain slices from each mouse. Student’s t-test, *** p-value < 0.001. NS, not significant. **(d)** Representative immuno-EM micrographs demonstrating the postsynaptic localization of anti-BirA-gold-silver particles in Adora2-cre mouse striatum injected with FLEX-postBirA* AAVs (left & middle). No specifically localized anti-BirA-gold-silver particle was observed in the other striatum without AAV injection from the same mouse (right). Black arrow = synaptic cleft particle, yellow arrow = intra spine particle. Scale bar, 500 nm. Sp, spine, AT, axonal terminal. **(e)** Quantification of synaptic clefts containing gold-silver particles (23 out of 43, 53%) and intra spine (20 out of 43, 47%). **(f)** Biochemical analyses using silver staining (top) and Strepavidin-HRP blot (bottom) confirms that the three BirA* probes are enzymatically active in Adora2-Cre mouse striatum. **(g)** Excitatory postsynaptic protein GluN2B, but not inhibitory postsynaptic protein GABA_A_1, is enriched in NeutrAvidin affinity purified material from postBirA* expressed striatum with biotin administration (top). Neither GluN2B nor GABA_A_1 were specifically enriched in the affinity purified materials from striatum expressing either of the other two BirA* probes (middle and bottom).

To use these tools to study cell-type specific synaptic proteomes, AAVs carrying the BirA* plasmids were stereotaxically injected into the striatum of Adora2-Cre mice, restricting their expression to A2a adenosine receptor expressing iSPNs ^11,12^. After one month, all three BirA* probes were robustly expressed **(Fig. 1a, top panel & Extended Data Fig. 1c)**. The postBirA* probe exhibited a punctate expression pattern in GFP-expressing iSPNs **(Fig. 1b & Extended Data Fig. 1d, left panels)**. In contrast, both the preBirA* and cytoBirA* probes had a diffuse expression pattern **(Fig. 1b & Extended Data Fig. 1d, middle & right panels)**. The postBirA* puncta co-localized with excitatory postsynaptic marker PSD95, with significantly less colocalization with the inhibitory postsynaptic marker gephyrin **(Fig. 1c)**. PostBirA* also was commonly juxtaposed with presynaptic vesicular glutamate transporter 1 (VGLuT1) but not the vesicular GABA transporter (VGAT) **(Extended Data Fig. 1e)**. Thus, the postBirA* probe was enriched at the postsynaptic membrane of excitatory SPN synapses. By contrast, neither preBirA* nor cytoBirA* colocalized with PSD95 or gephyrin **(Fig. 1c)**.

Next, we performed immuno-electron microscopy (iEM) to confirm postsynaptic localization of postBirA* in SPNs **(Fig. 1d-e & Extended Data Fig. 1f)**. FLEX-postBirA* AAVs were injected into one striatum of Adora2-Cre mice and the other striatum was used as a negative control for iEM. Anti-BirA-gold-silver particles were found either in the synaptic cleft or within spine heads. Very few gold-silver particles were detected, and none localized to synapses in the uninfected striatum **(Fig. 1d & Extended Data Fig. 1f, right panels)**. Our iEM data strongly suggest that, when expressed in striatal SPNs, the biotin ligase domain of postBirA* is present at similar levels within the postsynaptic density and in spine heads. Since BirA* selectively biotinylates proteins in close physical proximity ^17^, our results suggested that postBirA* would predominantly biotinylate a broad spectrum of postsynaptic proteins in SPN glutamatergic synapses, including both transmembrane (e.g., AMPARs) and scaffolding proteins (e.g., PSD95).

Next, we examined the proteins biotinylated by each of BirA* probes in Adora2-Cre mice following subcutaneous injection of biotin or vehicle (i.e., saline) **(Fig. 1f)**. Biotinylated proteins were affinity purified with NeutrAvidin agarose and the purified material was analyzed with SDS-PAGE and silver staining **(Fig. 1f, top panels)**. Streptavidin-horseradish peroxidase blot was used to detect biotinylated proteins **(Fig. 1f, bottom panels)**. Robust protein biotinylation signals were detected in the affinity-purified material from striata expressing the BirA* probes following biotin administration. For example, GluN2B, an NMDAR subunit known to be present at excitatory postsynaptic terminals, was only detected in the affinity-purified material from striata expressing postBirA* **(Fig. 1g, top panel)**. The strongest GluN2B signal was detected in samples expressing postBirA* with biotin administration, while GABA_A_1, a marker of inhibitory synapses was not enriched **(Fig. 1g, top panel)**. Importantly, nearly no GluN2B signal was detected in samples purified from striata expressing preBirA* or cytoBirA* **(Fig. 1g & Extended Data Fig. 1g)**. We then used IHC to examine where the biotinylated proteins were localized. Notably, we found more biotin puncta colocalized with PSD95 in striata expressing postBirA* than in striata expressing the other probes **(Extended Data Fig. 1h)**. In summary, these results show that postBirA* can selectively biotinylate postsynaptic proteins at glutamatergic synapses *in vivo*.

We next examined whether postBirA* affects synaptic function when expressed in mouse striatum. To this end, we injected Cre-independent channelrhodopsin (ChR2) into primary motor cortex M1 of Adora2-Cre mice and FLEX-postBirA* or FLEX-cytoBirA* into dorsolateral striatum (DLS) of the same mice **(Extended Data Fig. 2a-c)**. Five weeks later, *ex vivo* brain slices were prepared from these mice and optogenetic methods used to activate corticostriatal glutamatergic axons while patch clamp recording from iSPNs in the DLS. The amplitude of light-induced excitatory postsynaptic currents (EPSCs) in postBirA* and cytoBirA* expressing SPNs was very similar **(Extended Data Fig. 2d)**. To examine SPN GABAergic synaptic transmission, Cre-independent ChR2 was co-injected with preBirA* or cytoBirA* into the striatum of Adora2-Cre mice **(Extended Data Fig. 2e-f)**. This led to robust expression of ChR2 in SPNs projecting to the GPe. In *ex vivo* brain slices from these mice, optogenetic methods were used to activate striatopallidal axons while recording from GPe neurons using patch clamp methods. The amplitude of inhibitory postsynaptic currents (IPSC) recorded in GPe neurons from preBirA* and cytoBirA* injected mice were indistinguishable **(Extended Data Fig. 2g-h)**. These findings suggest that our BirA* probes do not alter iSPN pre- or post-synaptic function.

To quantitatively compare iSPN and dSPN postsynaptic compartment proteomes, we performed a postBirA*-based 10-plex TMT mass spectrometry (MS) experiment **(Fig. 2a)**. To minimize the impact of harvesting extraneous proteins ^18^, a reference library of postsynaptic terminal proteins was constructed by supervised machine learning **(Extended Data Fig. 3 & Table S1)**. Only proteins belonging to this library were considered relevant. As expected, the postsynaptic proteomes of iSPNs and dSPNs were similar **(Fig. 2b & c, Table S2)**. However, there were differences. For example, the catalytic subunit of protein kinase A (Prkaca) was more abundant in the iSPN proteome than that of dSPNs ^19,20^. This difference was confirmed by WB and IHC **(Extended Data Fig. 4a-e)**. Additional evidence that this difference was of functional significance came from the discovery that there was a higher level of serine/threonine phosphorylated synaptic proteins in iSPNs than dSPNS **(Extended Data Fig. 6 & Table S2)**.

**Fig. 2.**
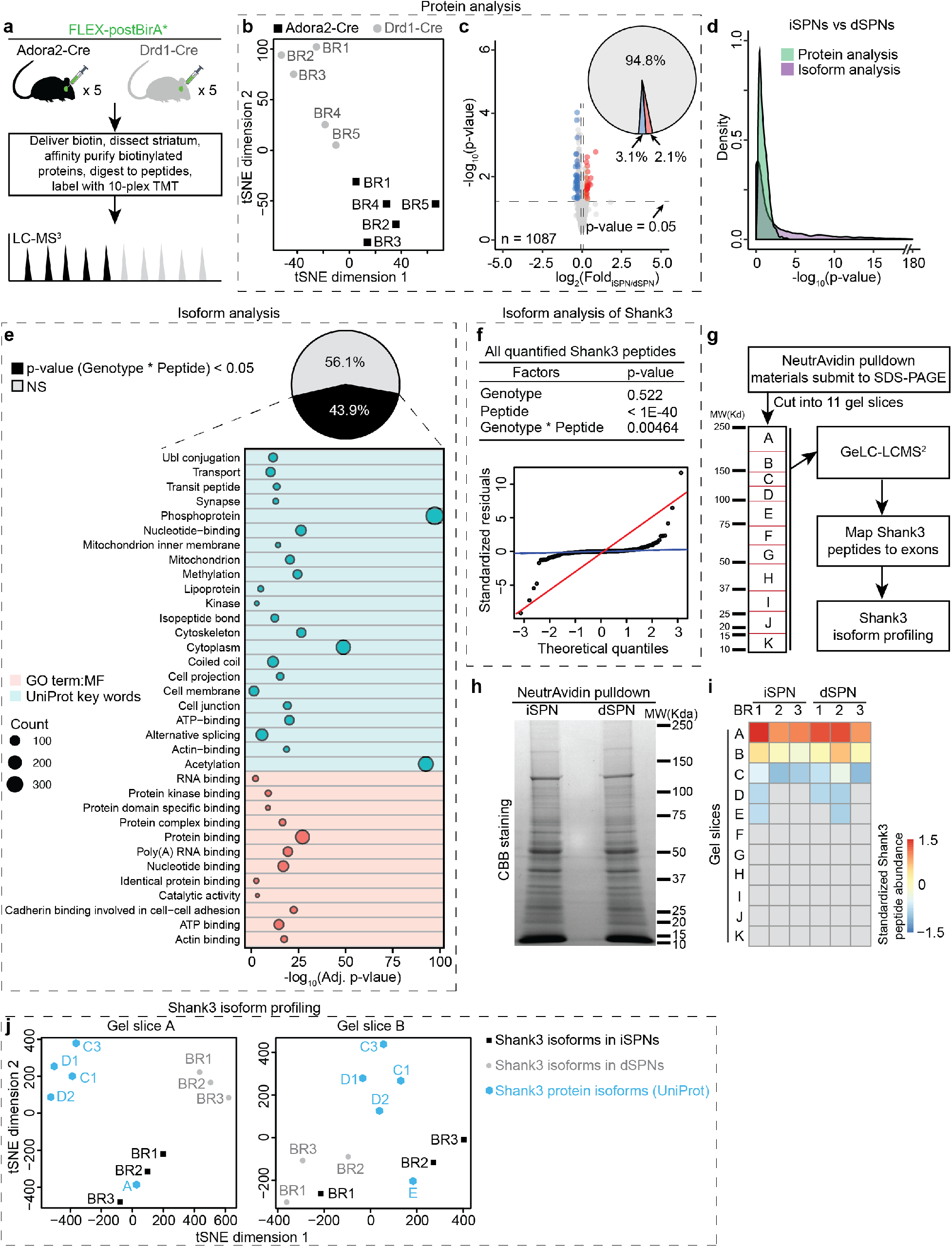
Differences between iSPN and dSPN postsynaptic compartment proteomes are predominantly at the isoform level. **(a)** Experimental design to compare iSPN and dSPN postsynaptic compartment proteomes. **(b)** Biological replicates (BR) cluster by genotype in (t-distributed stochastic neighbor embedding) tSNE plot based on protein analysis. **(c)** Volcano plot depicting comparison of iSPN and dSPN postsynaptic compartment proteomes based on protein analysis. Pie chart showing only ∼5% proteins in iSPN postsynaptic compartment proteomes are differentially expressed in dSPNs. For proteins with multiple UniProt accessions, we only considered the accession with highest total TMT reporter ion intensity. **(d)** Isoform analysis revealed that the postsynaptic proteins were divergently expressed in iSPNs and dSPNs at isoform level. For protein analysis, p-values were calculated by Student’s t-test. For isoform analysis, p-values were calculated by two-way ANOVA. (PR > F): Genotype * peptide values were used. **(e)** Gene functional classification analysis of postsynaptic proteins isoform-divergently expressed in iSPNs and dSPNs. For all listed terms, count > 30, adj. p-value < 0.05. **(f)** Top, p-values acquired by two-way ANOVA analysis of Shank3 peptides. Bottom, quantile-quantile (Q-Q) plots showing that the variances of Shank3 peptides in iSPNs and dSPNs are heteroscedastic. **(g)** Experimental design of GeLC-LCMS^2^ based Shank3 protein isoform profiling. **(h)** Representative Coomassie blue (CBB) staining gels of NeutrAvidin affinity purified material from iSPN and dSPN samples. **(i)** Heatmap showing the distribution pattern of all identified Shank3 peptides in gel slices. **(j)** Profiling of Shank3 protein isoforms in iSPNs and dSPNs postsynaptic compartments. Left, in gel slice A, the Shank3 protein isoform(s) expressed in iSPNs were closely-clustered with Shank3A. In contrast, the isoform(s) expressed in dSPNs were separated from others, suggesting there is unreported long Shank3 isoform(s) co-expressed with Shank3A in dSPNs. Right, in gel slice B, the isoforms expressed in dSPNs and iSPNs were loosely clustered with Shank3E.

Since ∼70% of mammalian genes can produce multiple protein isoforms ^21,22^, we hypothesized that traditional protein analysis workflow is not well suited to identify dissimilarities of SPNs at the protein isoform level **(Extended Data Fig. 5a)**. Therefore, a two-way ANOVA-based isoform analysis was used to compare iSPN and dSPN proteomes **(Fig. 2d, Table S2)**. Surprisingly, nearly half (44%) of the synaptic proteomes of iSPNs and dSPNs differed at the isoform level **(Fig. 2d & Extended Data Fig. 5b-c)**. As different protein isoforms can have different functions ^21^, the disparity between iSPNs and dSPNs could have physiological consequences.

One of the most prominent differences between SPNs at the isoform level was for the postsynaptic scaffolding protein Shank3 ^23-25^ **(Fig. 2f)**. Shank3 physically interconnects glutamate receptors with PSD-95 and Homer to the actin cytoskeleton and choreographs dendritic spine and synapse formation, maturation, maintenance, and plasticity. Shank3 gene mutations cause several of neuronal developmental disorders, such as ASD, Phelan-McDermid syndrome (PMS), schizophrenia and intellectual disability (ID) ^23,24^. To validate this finding, we performed in-gel digestion coupled with mass spectrometric analysis (GeLC-MS2) to profile Shank3 protein isoforms in iSPNs and dSPNs **(Fig. 2g-i & Extended Data Fig. 7a)**. By denominational analysis of the identified Shank3 peptides in each gel piece, we found that the major long Shank3 isoforms expressed in iSPN and dSPNs postsynaptic proteomes differed **(Fig. 2j & Extended Data Fig. 7b-c, Table S3)**. The predominant long Shank3 protein isoform expressed in iSPNs and dSPNs was similar to Shank3A, but dSPNs also expressed another isoform(s). In contrast, iSPNs and dSPNs expressed similar short Shank3 protein isoforms.

To explore the potential functional significance of this difference, *Shank3B*^*-/-*^ mice were examined. *Shank3B*^*-/-*^ mice have significant postsynaptic impairments in the striatum and display robust ASD-like behaviors, such as self-injurious grooming ^26^. Therefore, our findings using this model should help guide the interpretation *Shank3B*^*-/-*^ mouse phenotypes and may provide insight into human neurological conditions stemming from *Shank3 gene* mutations. In this mouse line, *Shank3* gene exons 13-16 have been replaced with a neomycin resistance cassette **(Extended Data Fig. 8a)**. Full-length Shank3A protein requires all 22 exons, while Shank3E protein is encoded by exons 17-22 ^24,27^. Thus, in *Shank3B*^*-/-*^ mice, Shank3A was completely depleted while Shank3E remained ^26^. Although Shank3C and D were also disrupted in this mouse line, their expression in striatum is very low and is not expected to play a major role in striatal synaptic function ^23,24,28^. Two independent postBirA*-based 10-plex TMT MS experiments were performed to compare postsynaptic alterations in *Shank3B*^*-/-*^ iSPNs and dSPNs **(Fig. 3a)**. This analysis revealed that the iSPN postsynaptic proteome was more profoundly altered in *Shank3B*^*-/-*^ mice than was that of dSPNs **(Fig. 2b-c, Table S4)**. We further confirmed divergently regulated proteins identified through *in vivo* proximity biotin-tagging TMT experiments by WB **(Fig. 3d-e)**. These results confirmed that the glutamatergic synaptic proteome of iSPNs was more dramatically altered than that of dSPNs **(Fig. 3f-h & Extended Data Fig. 8b, Table S4)**. These findings suggest that although Shank3A is a major postsynaptic scaffold in both iSPNs and dSPNs, dSPNs express another Shank3 isoform(s) that can partially compensate for the loss of Shank3A.

**Fig. 3.**
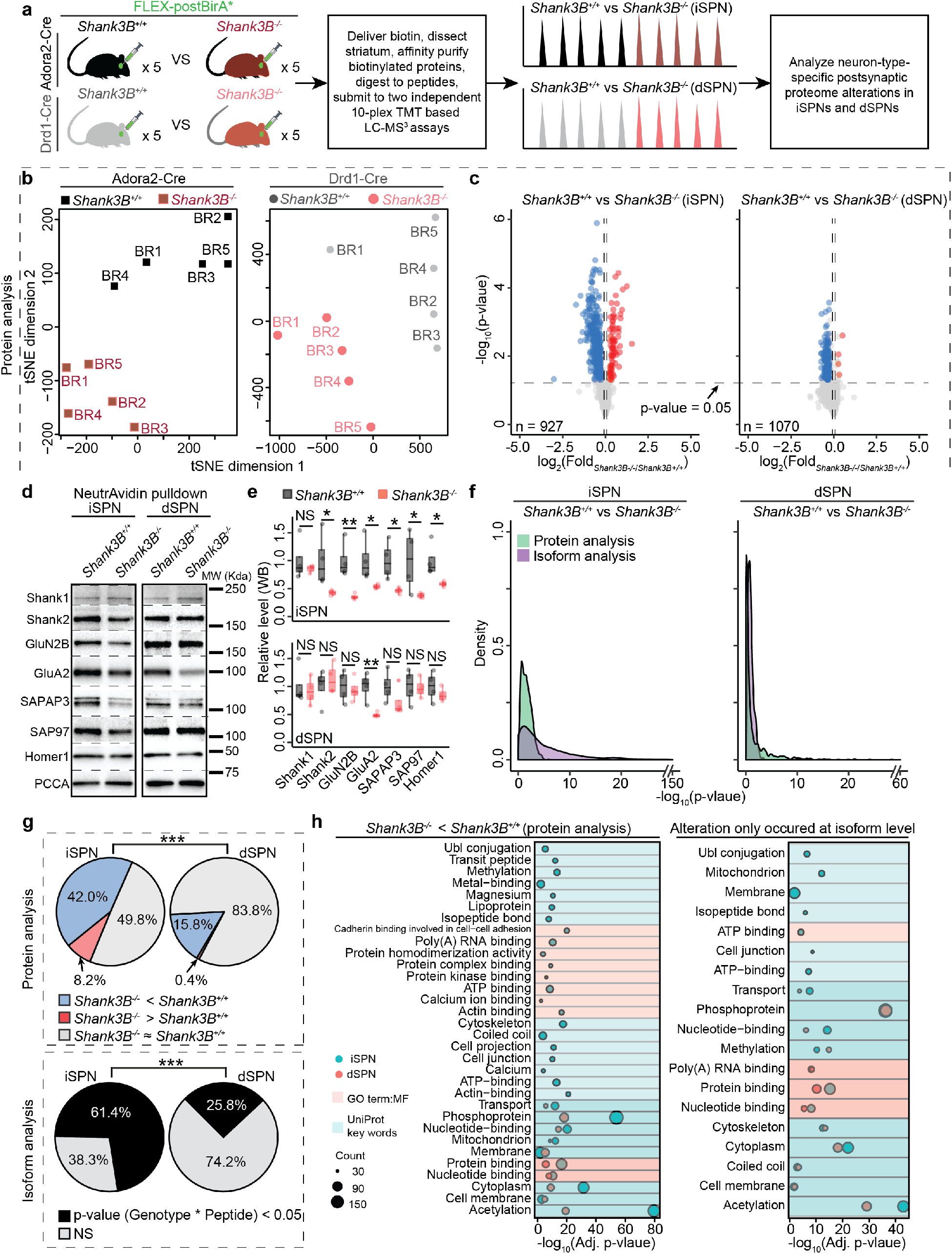
Deletion of *Shank3* gene exons 13-16 divergently altered iSPN and dSPN postsynaptic compartment proteomes. **(a)** Experimental design to profile iSPN and dSPN postsynaptic compartment proteome alterations in *Shank3B*^*-/-*^ mice. **(b)** Biological replicates cluster by genotype in tSNE plots based on protein analysis. **(c)** Protein analysis revealed that ISPN postsynaptic compartment proteome was altered to a greater degree in *Shank3B*^*-/-*^ mice compared to dSPNs. For proteins with multiple UniProt accessions, we only considered the accession with highest TMT reporter ion intensity. **(d)** SPN-type specific alterations of some proteins (Shank2, GluN2B, SAPAP3, SAP97 and Homer1) were validated by WB. GluA2 level was significantly reduced in both *Shank3B*^*-/-*^ iSPNs and dSPNs compared to *Shank3B*^*+/+*^ iSPNs. Shank3B deletion didn’t alter Shank1 level in SPNs. **(e)** Quantification of **(d)**, protein levels are normalized to PCCA. n = 4 mice per genotype. **(f)** Exons 13-16 deletion led to a more robust iSPN postsynaptic protein expression alteration at isoform level compared to dSPN. **(g)** Comparisons of postsynaptic protein alterations in *Shank3B*^*-/-*^ iSPNs and dSPNs revealed by protein analysis and isoform analysis. **(h)** Left, gene functional classification analysis of postsynaptic proteins significantly downregulated in *Shank3B*^*-/-*^ iSPNs and dSPNs (protein analysis). Right, gene functional classification analysis of postsynaptic proteins which were differentially expressed (i.e. significantly) at isoform level in *Shank3B*^*+/+*^ and *Shank3B*^*-/-*^ SPNs. **(e)** Student’s t-test, **(g)** Fisher’s exact test. * p-value < 0.05, ** p-value < 0.01, *** p-value < 0.001, NS, not significant.

To test this hypothesis, we performed an additional isoform analysis of the *Shank3B*^*-/-*^ postsynaptic proteomes. The relative abundance of Shank3 peptides was homoscedastic in *Shank3B*^*-/-*^ iSPNs, but was highly heteroscedastic in *Shank3B*^*-/-*^ dSPNs **(Fig. 4a & Extended Data Fig. 8c)**. This result strongly suggests that the Shank3 protein isoforms in *Shank3B*^*-/-*^ iSPNs and dSPNs were dissimilar. We further performed Shank3 WB analysis of NeutrAvidin agarose-affinity purified material from postBirA*-expressed striata **(Fig. 4b)**. Interestingly, there was very little Shank3 signal in *Shank3B*^*-/-*^ iSPNs. However, in *Shank3B*^*-/-*^ dSPNs, several Shank3 protein isoforms were present. As there is not an antibody-based strategy that is able to detect all Shank3 protein isoforms, we performed GeLC-MS2 to profile the Shank3 protein isoforms in *Shank3B*^*-/-*^ SPNs **(Fig. 4c-e & Extended Data Fig. 8d, Table S5)**. Shank3E was still expressed in *Shank3B*^*-/-*^ dSPNs. We also identified two unreported Shank3 isoforms in *Shank3B*^*-/-*^ dSPNs.

**Fig. 4.**
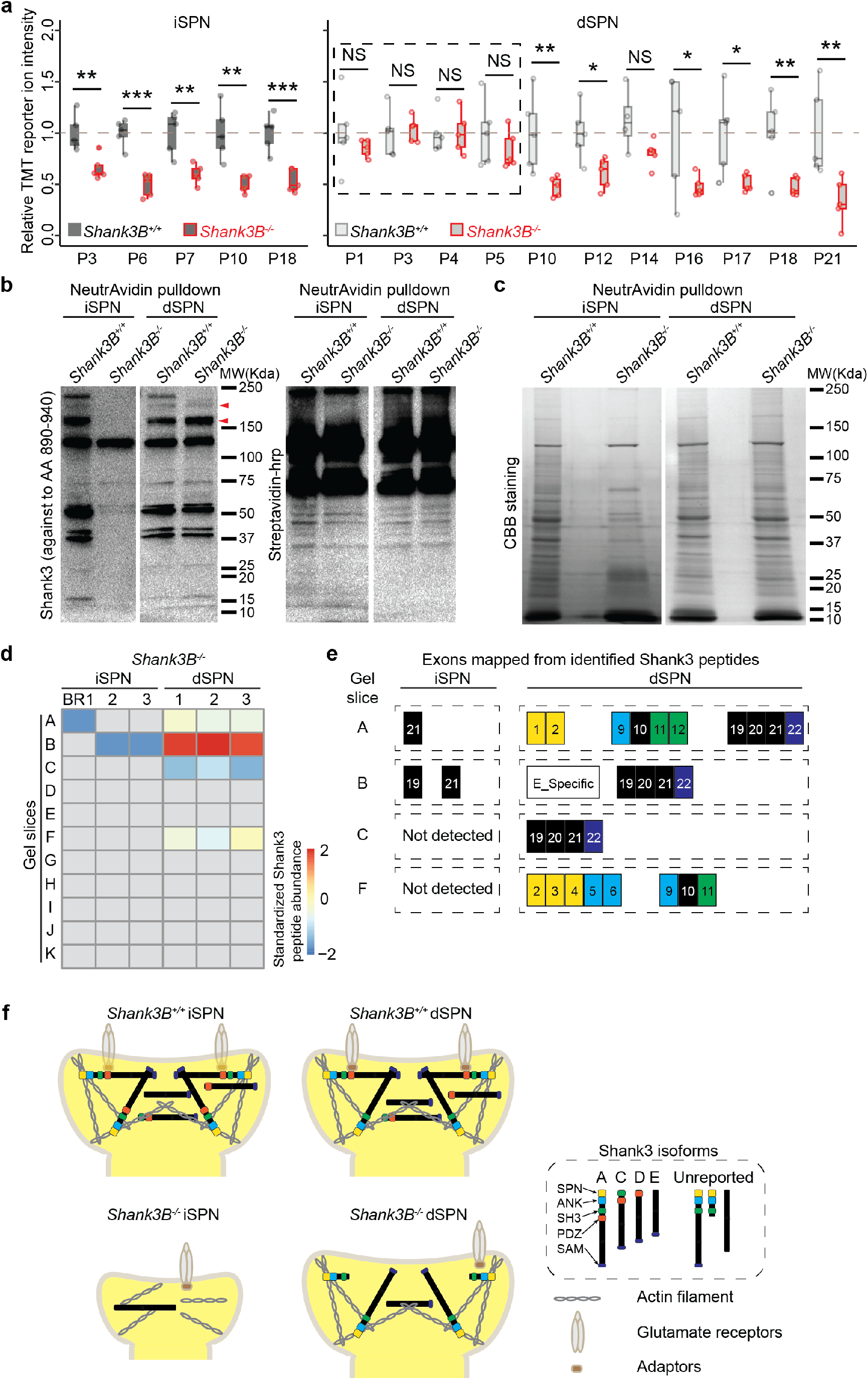
Distinct Shank3 protein isoforms were expressed in dSPNs and iSPNs postsynaptic compartments. **(a)** The relative abundances of Shank3 peptides were homoscedastic in *Shank3B*^*-/-*^ iSPNs but heteroscedastic in *Shank3B*^*-/-*^ dSPNs. P = quantified Shank3 peptide. **(b)** Left, representative WB anlaysis of NeutrAvidin affinity purified material showed dissimilar Shank3 protein isoform expression patterns in *Shank3*^*-/-*^ SPNs. Three bands between 150-250 Kd in *Shank3B*^*+/+*^ SPN samples are usually considered as Shank3A, 3C/D and 3E. Arrow heads indicated Shank3 isoforms expressed in *Shank3*^*-/-*^ dSPN sample. Right, representative blots showed the level of all biotinylated protein as loading control. **(c)** Representative CBB staining gels of NeutrAvidin affinity purified material from the indicated SPN samples. **(d)** Shank3 peptides were identified at multiple gel slices. In *Shank3*^*-/-*^ iSPNs, we only identified a few Shank3 peptides in gel slices A and B. However, in *Shank3*^*-/-*^ dSPNs, Shank3 peptides were detected in gel slices A, B, C & F. **(e)** Identified Shank3 peptide amino acids were mapped to in silico translated *Shank3* gene exons (NM_021423.4). Each colored block represents a *Shank3* gene exon mapped from the identified peptides. Numbers in blocks, exon numbers. E_Specific, this peptide only belongs to Shank3E. Yellow, SPN domain, sky blue, ANK, green, SH3, dark blue, SAM. The data suggest that there may be an unreported long Shank3 isoform(s) in dSPN postsynaptic compartment which do not contain the PDZ domain. Notably, there we detected an unreported short Shank3 isoform in *Shank3*^*-/-*^ dSPN postsynaptic compartments, which contains SPN, ANK and SH3 domains. **(f)** Working model of the molecular mechanism underlying SPN-type-specific postsynaptic compartment impairments in *Shank3*^*-/-*^ striatum. ANK, ankyrin repeats domain, SH3, src 3 domain, SAM, sterile α motif domain, SPN, N-terminal domain.

Moreover, the Shank3 proteins isoforms expressed in *Shank3B*^*-/-*^ dSPNs contained all of the protein domains for cytoskeletal organization and Homer binding ^24^ **(Fig. 4e)**. Therefore, these isoforms could still serve as postsynaptic scaffoldings to maintain the normal morphology of dendritic spines in *Shank3B*^*-/-*^ dSPNs. Our findings highlight that cell-type-specific Shank3 protein expression leads to a divergence in the impact of the *Shank3B*^*-/-*^ deletion on iSPNs and dSPNs **(Fig. 4f)**.

Patients with *Shank3* gene mutations display a wide range of clinical symptoms ^23,24,29^. For example, PMS patients with only have one *Shank3* gene copy, suffer from delayed development, hypotonia, delayed or absent speech, and autistic symptoms such as restricted and repetitive behaviors. Other *Shank3* gene mutations, such as non-sense mutations are frequently observed in ASD and ID patients. Similarly, ASD mouse models with Shank3 gene mutations manifest some ASD-like deficits, but also have a wide range of neurological phenotypes. This phenotypic heterogeneity has been hypothesized to be due to the expression pattern of Shank3 protein isoforms and their complex impact on postsynaptic proteomes ^28,30^. Consistent with this thesis, we found clear differences between iSPNs and dSPNs in their expression of Shank3 protein isoforms. These results are consistent with recent work showing that the synaptic impairment in *Shank3B*^*-/-*^ mice differs between iSPNs and dSPNs ^31^, which may disrupt striatal motor control and cause repetitive behaviors.

Many other synaptic genes associated with neurodevelopmental disorder, such as *Syngap1* ^32^, *Kalrn* ^33^, *Nlgns*, and *Nrxns* ^34^ are also expressed in multiple isoforms. Therefore, the pathological mutation of these risk genes may also lead to protein isoform imbalances and postsynaptic proteome alterations in a neuron-type-specific manner. Our methodology can be used to examine neuron-type-specific protein expression patterns and direct pharmaco- and genetic therapies for treating neuronal disorders, such as ASD.

## Methods

All materials used in this study are available upon request.

### Animals

All procedures were approved by Northwestern University’s Animal Care and Use Committee (IS00001789, IS00010673, IS00009321) in compliance with US National Institutes of Health standards. *Shank3B*^*-/-*^ mice (B6.129-Shank3tm2Gfng/J, Stock No: 017688) were acquired from The Jackson Laboratory. We acquired frozen sperm of Drd1-Cre mouse (EY262Gsat/Mmucd, GENSAT Project, backcrossed to C57BL/6J background, MMRRC ID, 17264) from Mutant Mouse Resource & Research Centers and bred Drd1-Cre mice with the help from Northwestern university Center for Comparative Medicine. Adora2-Cre mice, KG139Gsat/Mmucd, GENSAT Project, backcrossed to C57BL/6J background, MMRRC ID, 36158.

We first crossed homozygous Drd1-Cre or Adora2-Cre mice with *Shank3B*^*-/+*^ mice to achieve Drd1-Cre^-/+^::*Shank3B*^*-/+*^ and Adora2-Cre^-/+^::*Shank3B*^*-/+*^ mice. Then we bred Drd1-Cre^+/+^::*Shank3B*^*-/+*^ and Adora2-Cre^+/+^::*Shank3B*^*-/+*^ and backcrossed the mice to obtain Drd1-Cre::*Shank3B*^*+/+*^, Drd1-Cre::*Shank3B*^*-/-*^, Adora2-Cre::*Shank3B*^*+/+*^ and Adora2-Cre::*Shank3B*^*-/-*^ mice. For all TMT-MS experiments, eight-week old mice were used for intracranial AAV injections. 10-plex TMT-MS experiments compared postsynaptic compartment proteomes of *Shank3B*^*+/+*^ and *Shank3B*^*-/-*^ SPNs littermate controls. For biochemical and IHC experiments, 8-12 weeks old mice were used. For biochemical, IHC and 3-plex TMT-MS experiments, mice were randomly selected. Both male and female mice were used for all experiments.

### Cloning

Our probes were based on BirA*-based proximity biotin-tagging ^17^ and the synaptic localization motifs from mGRASP ^35^. We obtained pcDNA3.1 MCS-BirA(R118G)-HA, paavCAG-pre-mGRASP-mCerulean and paavCAG-post-mGRASP-2A-dTomato from Addgene (Cat #, 53581, 34910 and 34912, RRID: Addgene_53581, Addgene_34910 and Addgene_34912). Overlap extension PCR was used to construct BirA* probes **(Fig. S1A)**. For postBirA*, we individually amplified SP, BirA*-HA, tr-mNL1 and 2A-GFP fragments by PCR. Then we constructed SP-BirA*-HA and tr-mNL1-2A-GFP by linking corresponding fragments using overlap extension PCR. These two fragments were digested by EcoRI (New England Biolabs, Cat # R0101) and then linked together using T4 DNA ligase (New England Biolabs, Cat # M0202) to form postBirA*. Then postBirA* fragment and pAAV-FLEX-GFP vector (Addgene, Cat # 28304) were both digested with KpnI (New England Biolabs, Cat # R3142) and BsrgI (New England Biolabs, Cat # R0575). FLEX-postBirA* was formed by linking digested postBirA* fragment with linearized pAAV-FLEX-GFP vector **(Fig. S1B)**. Thus, the GFP coding sequence was replaced to postBirA*. FLEX-cytoBirA* and FLEX-preBirA* were constructed in a similar way.

### Stereotaxic delivery of AAVs

Adora2-Cre::*Shank3B*^*+/+*^, Adora2-Cre::*Shank3B*^*-/-*^, Drd1-Cre::*Shank3B*^*+/+*^ or Drd1-Cre::*Shank3B*^*-/-*^ mice were anesthetized with isoflurane (Covertrus, Cat # 029405) before being placed in a stereotaxic injection rig (KOPF, model 922). Mouse head was restrained by ear bars with placing the front teeth in the holding apparatus. Isoflurane and oxygen were provided (Kent, VetFlo) at a low flow rate to keep the mouse anesthetized during the whole procedure. The animal’s body temperature was maintained at 37–38°C with an electric heating pad (Homeothermic blanket system, Harvard Apparatus). To prevent from drying, the eyeballs were covered with ophthalmic ointment (Artificial tears, Henry Schein). Buprenex (0.1 mg/kg) was administrated by intraperitoneal (IP) injection. AAVs were injected at two positions in each striatum using Hamilton needles (Hamilton, 65460-02). Injection sites, ±2.4 mediolateral, 0.5 anteroposterior, –3.0 and –3.5 dorsoventral. An automatic injection pump (World Precision Instruments) was used to control the speed at 200 nl/min. The needle was holding at injection position for 5 mins before slow withdrawal. After closing the wound by wound clips, meloxicam (1 mg/kg) was administrated by IP injection. The mouse was put onto a warming pad for recovery. The next morning, meloxicam was administrated one more time to the AVV-infected mouse. One month later, biotin (24 mg/kg in saline) was administrated by subcutaneous (SubQ) injection for seven consecutive days ^36^. For the no-biotin control experiments in **Fig.1F-G**, saline was administrated by SubQ injection for seven consecutive days.

### Electrophysiology and data analysis

Experiments were performed using acute brain slices from 4-5-month-old Adora2-Cre mice. The brains were rapidly removed and cooled in ice-cold oxygenated sucrose-Artificial cerebrospinal fluid (ACSF), composed of (in mM): Sucrose 220; KCl 2.5; CaCl_2_ 0.5, MgSO_4_ 3, NaH_2_PO_4_ 1.2, NaHCO_3_ 26, glucose 5. Coronal and para-sagittal slices (300 μm) were prepared in ice-cold oxygenated (95% O_2_ / 5% CO_2_) ACSF with a vibratome (LeicaVT1000S, Leica Biosystems, Germany), then warmed to 36°C for 45 mins, allowed to cool to room temperature, and transferred as needed to a submerged slice chamber mounted on the stage of an upright microscope, perfused at 2 ml/min with oxygenated normal ACSF, comprised of (in mM): NaCl 124; KCl 3.5, CaCl_2_ 2.5, MgSO_4_ 1.2, NaH_2_PO_4_ 1.2, NaHCO_3_ 26, glucose 11. Cell attach and whole-cell patch clamp recordings in voltage clamp mode were made using a MultiClamp 700B amplifier (Molecular Devices, California, Sunnyvale, USA. Recordings were performed with patch pipettes (3-5 MΩ resistance) containing (in mM): K-gluconate 145, EGTA 1.1, HEPES 10, CaCl_2_ 0.1, MgCl_2_ 4, Na_2_ATP 2, NaGTP 0.3, pH adjusted to 7.2. EPSCs were recorded from indirect pathway spiny projection neurons (iSPNs) in voltage-clamp mode at a holding potential of -80 mV. IPSCs were recorded from globus pallidus (GP) neurons in voltage-clamp mode at a holding potential of -40 mV. Evoked responses for input-output curves were averaged from n = 3 per stimulus intensity. Both input-output curves were derived from the peaks of the evoked currents.

Stereotaxic injections: Stereotaxic injections were performed using a computer-assisted stereotaxic system (Leica Biosystems, Buffalo Grove, IL). Mice were anesthetized with isoflurane. The injection coordinates were: for striatal injections of cytoBirA*, preBirA* and pAAV-EF1a-double floxed-hChR2(H134R)-mCherry-WPRE-HGHpA (lateral, 2.70 mm; posterior, -0.10 mm; depth, 3.40 mm); for dorsolateral striatum (DLS) injections of cytoBirA* and postBirA* (lateral, 2.15 mm; posterior, 1.18 mm; depth, 1.33 mm), and for the cortical injections of pAAV-hSyn-hChR2(H134R)-mCherry (lateral, 1.51 mm; posterior, 0.98 mm; depth, 3.40 mm). To maximize the extent of the infection, the AAV-containing solution was injected at a moderate speed (6-8 psi, 60-100 secs duration) (IM 300 Microinjector, Narishige, Japan). Experiments with Cre-on construct were performed 4 weeks after injection whereas experiments with a ChR2 Cre-independent construct were performed two weeks after virus injection.

Data analysis: Evoked EPSCs and IPSCs were analyzed with pClamp 10 (Molecular Devices, California, Sunnyvale, USA) program. Statistical analysis was performed using GraphPad Prism 6.07 (GraphPad Software, Inc., La Jolla, CA, USA) and included either a one-way or a two-way ordinary ANOVA, followed by Bonferroni’s multiple or comparisons test. Data are presented as mean ± SEM.

### Immunohistochemistry (IHC) staining

Para-sagittal brain slices (200 μm) were prepared as descried above. Slices were briefly fixed in 4% paraformaldehyde (PFA, VWR, Cat # AA-A11313-36) in phosphate buffered saline (PBS, pH 7.4) for 15 mins with mild agitation at room temperature and then washed 3 × 5 mins in glycine solution (1M glycine in PBS, pH 7.4) to block unreacted PFA. Slices were further washed 3 × 5 mins in PBS to remove glycine. For permeabilization, slices were incubated in Triton X-100 (0.5 % in PBS, pH7.4) solution for 1 hr. After blocking (10% horse serum, 0.5 % Trition X-100 in PBS, pH7.4) for 1 hr, slides were incubated with primary antibodies at 4°C for 48 hrs with mild agitation. Following primary antibodies were used: goat anti-BirA (1:200, MyBioSource, Cat # MBS534316, RRID:AB_10578606), chicken anti-GFP (1:1000, Abcam, Cat # ab13970, RRID:AB_300798), guinea pig anti-VGluT1 (1:1000, Millipore Sigma, Cat # AB5905, RRID:AB_2301751), rabbit anti-Gephyrin (1:500, Synaptic System, Cat # 147008, RRID:AB_2619834), mouse anti-PSD95 (1:400, Thermo Fisher Scientific, Cat # MA1-046, RRID:AB_2092361), rabbit anti-VGAT (1:500, Synaptic System, Cat # 131003, RRID:AB_887869) and rabbit anti-Prkaca (1:500, Millipore Sigma, Cat # HPA071185, RRID:AB_2686356). After 6 × 5 min washing in PBS, slides were incubated with corresponding Alexa Fluor secondary antibodies (1:1000, Thermo Scientific, Cat #, A11030, A11057, A21057, A31571, A31573 and SA5-10098. RRID: AB_2534089, AB_2534104, AB_2535723, AB_162542, AB_2536183 and AB_2556678) or NeutrAvidin-Rhoaminne Red (1:2000, Thermo Fisher Scientific, Cat # 6378) overnight at 4°C with mild agitation. Then, after 3 × 5 mins washing in PBS, slides were incubated in 4′,6-diamidino-2-phenylindole (1:1000, DAPI, Sigma-Aldrich, Cat # D9542) solution for 10 mins. The slides were washed 3 × 5 mins and mounted with Fluoromount-G (SouthernBiotech, Cat # 0100-01) on microscope slides with Secure-Seal spacer (Invitrogen, Cat # S24736).

For IHC experiments to detect Prkaca expression in iSPNs and dSPNs (**Fig. S4E**), Drd1-Cre or A2a-Cre mice were anesthetized and transcardially perfused with ice old PBS followed by 4% PFA. Dissected brains were cryoprotected in 30% sucrose solution and then buried in embedding medium (Sakura, Cat# 4583). Coronal sections (35 μm) were obtained using a cryostat microtome. IHC staining procedure was the same as staining 200 μm thick brain slices. Rabbit anti-Prkaca (1:400, Millipore Sigma, Cat# HPA071185) was used to probe Prkaca expressions in SPNs.

Wole brain slice images were captured using TissueGnostics with 20× objective lens. Images of synaptic proteins were captured using a Nikon A1R+ confocal laser microscope and 60× or 100× objective lens. For each mouse, brain slices were randomly selected. Images were processed and analyzed by Fiji (NIH) with plugins Ratioplus, Colocalization Finder, and JACOP.

### Immunogold staining and silver enhancement

Para-sagittal brain slices (200 μm) were prepared as descried above. Slices were incubated with 4% PFA and 0.2% glutaraldehyde (Polysciences, Cat # BLI1909-10) in PBS (Corning, Cat #, 21-040-CV) for 15 mins followed by two washes with 0.1% NaBH_4_ (Sigma-Aldrich, Cat # 480886) in PBS (15 mins each). Then slices were washed by 3 × 15 mins with PBS and permeabilized with 0.005% Triton-X-100 (Sigma-Aldrich, Cat # X100) in PBS for one hours. Slices were incubated with Aurion blocking solution for one hour and washed 2 × 15 mins with incubation solution (Electron microscopy sciences, Cat # 25558). Then the slices were incubated with goat anti-BirA in incubation solution (1:50, MyBioSource, Cat # MBS534316, RRID:AB_10578606) at 4°C for 48 hrs with mild agitation. After 6 × 10 mins washed with incubation solution, sliced were incubated with donkey-anti-goat UltraSmall gold (1:50, Electron microscopy sciences, Cat #25800, RRID:AB_2631210) at 4°C for 24 hrs with mild agitation. Then slides were washed 6 × 10 mins with incubation solution and followed by 2 × 10 mins washes with PBS. Slices were post-fixed with 0.2% glutaraldehyde in PBS for 15 mins and washed 4 × 10 mins with MilliQ water. Slices were silver-enhanced with AURION R-GENT SE-EM kit (Electron microscopy sciences, Cat # 500.033) for 25 mins. Then slices were washed 3 × 10 mins in MilliQ water and sent to the Northwestern University Center for Advanced Microscopy for further processing. Briefly, samples were fixed in mixture of 2.5% glutaraldehyde and 2% paraformaldehyde in 0.1M cacodylate buffer for 2 or 3 hrs or overnight at 4°C. After post-fixation in 1% osmium tetroxide and 3% uranyl acetate cells were dehydrated in series of ethanol, embedded in Epon resin and polymerized for 48 hrs at 60°C. Then ultrathin sections were made using Ultracut UC7 Ultramicrotome (Leica Microsystems) and contrasted with 3% uranyl acetate and Reynolds’s lead citrate. Samples were imaged using a FEI Tecnai Spirit G2 transmission electron microscope (FEI Company, Hillsboro, OR) operated at 80 kV. Images were captured by Eagle 4k HR 200kV CCD camera. Images were processed and analyzed by Fiji (NIH).

### Affinity purification of biotinylated proteins

Dissected striata were homogenized in RIPA lysis buffer (50 mM Tris, 150 mM NaCl, 0.1% SDS, 1mM EDTA, 0.5% sodium deoxycholate, 1% Triton X-100, 1 x protease inhibitor cocktail (Thermo Fisher Scientific, Cat # 78443), 1 x phosphatase inhibitor (Thermo Fisher Scientific, Cat # 78420), pH 7.4) with an electronic homogenizer (Glas-Col, Cat # 099C-K54). Then excess 10% SDS solution was added into each sample to make the final SDS concentration to 1%. After sonication with a probe sonicator (Qsonica) for 3 × 1 min, striatal homogenates were solubilized at 4°C for one hr with rotation. Insoluble components were removed by centrifuging at 13,000 x g for 30 mins. 400 μl of pre-washed NeutrAvidin beads (Thermo Fisher Scientific, Cat # 2901) were added into each sample and incubated at 4°C overnight with gentle rotation.

For silver staining, Coomassie blue (CBB) staining and Western blotting (WB), after overnight incubation with striatal homogenates, NeutrAvidin beads were rinsed for 5 × 5 mins in 1 ml RIPA lysis buffer. Then 400 μl 2 x loading buffer were added into the beads. Mixtures were boiled at 95 °C for 5 mins and immediately put onto ice to elute the biotinylated proteins from the beads. Supernatants were carefully collected and reduced to ∼60 μl using a SpeedVac Vacuum Concentrator. Then all samples were brought up to 120 μl with RIPA lysis buffer. For silver staining, 20 μl of each sample was loaded into 4-12% Bis-Tris gels (Thermo Fisher, Cat # np0335box) for electrophoresis. For CBB staining and WB, 4-12% Tris-glycine gels (Invitrogen, Cat # XP04120BOX) were used.

### Silver and CBB staining

We used Pierce Silver Stain Kit (Thermo Fisher, Cat # 24612) for silver staining. Gels were washed 2 × 5 mins in ultrapure water and then fixed with 2 × 15 mins in 30% ethanol: 10% acetic acid solution. Then the fixed gels were washed 2 × 5 mins with 10% ethanol and 2 × 5 mins in ultrapure water. Then the gels were sensitized for 1 min and washed twice 2 × 1 min with water. Then, the gels were stained for 30 mins and washed 2 × 20 secs with ultrapure water. The gels were developed for 1-3 mins until bands appear and stopped with 5% acetic acid for 10 mins.

We used SimplyBlue™ SafeStain kit (Novex, Cat # LC6065) for CBB staining. Gels were washed 3 × 5 mins in ultrapure water and stained with SimplyBlue™ SafeStain solution overnight at room temperature (RT) with mild agitation. The gels were washed for 2 × 1 hour in MiiliQ water to reduce background staining.

### Western blotting (WB) and Streptavidin-horseradish peroxidase (HRP) blotting

After electrophoresis, proteins were transferred onto nitrocellulose membranes (Thermo Fisher, Cat # 4500002). For WB, the membranes were blocked with 10% bovine serum albumin (BSA, Jackson immunoResearch laboratories, Cat # 001-000-162) in TBST solution (Tris-buffered saline, 0.1% Tween 20) for one hour in RT. Then membranes were incubated with primary antibodies in incubation solution (3% BSA in TBST) overnight at 4°C with mild agitation.

Following primary antibodies were used: mouse anti-HA (1:1000, BioLegend, Cat # 901502, RRID:AB_2565007), chicken anti-GFP (1:2000, Abcam, Cat # ab13970, RRID:AB_300798), rabbit anti-PCCA (1:1000, NOVUSBIO, Cat # NBP2-32215), rabbit anti-GABAA1 (1:1000, EMD Millipore, Cat # 06-868, RRID:AB_310272), mouse anti-GluN2B (1:1000, Millipore Sigma, Cat # 05-920, RRID:AB_417391), rabbit anti-Prkaca (1:1000, Millipore Sigma, Cat # HPA071185, RRID:AB_2686356), rabbit anti-GluA2 (1:1000, Abcam, Cat # ab133477, RRID:AB_2620181), rabbit anti-SAPAP3 (1:1000, ThermoFisher scientific, Cat # 55056-1-AP, RRID:AB_10858793), rabbit anti-Homer1 (1:1000, Synaptic System, Cat # 160003, RRID:AB_887730), rabbit anti-SAP97 (1:1000, ThermoFisher scientific, Cat # PA1-741, RRID:AB_2092020), rabbit anti-GAPDH (1:2000, Cell Signaling Technology, Cat # 2118, RRID:AB_561053), rabbit anti-Shank1 (1:1000, Synaptic System, Cat # 162002), rabbit anti-Shank2 (1:1000, Cell Signaling Technology, Cat # 12218, RRID:AB_2797848), rabbit anti-Shank3 (1:1000, Boster Bio, Cat # A01231-1). After 3 × 10 min intense wash in TBST solution, membranes were incubated with corresponding secondary antibodies for one hour at RT with mild agitation. Following secondary antibodies were used: goat-anti-rabbit poly-HRP (1:2000, Invitrogen, Cat # 32260, RRID:AB_1965959), goat-anti-mouse poly-HRP (1:2000, Invitrogen, Cat # 32230, RRID:AB_1965958) and goat-anti-chicken HRP (1:2000, Abcam, Cat # ab97135, RRID:AB_10680105). Then membranes were washed 3 × 10 mins in TBST and developed with SuperSignal West Pico Chemiluminescent Substrate (Thermo Fisher, Cat# 34578) and imaged on a Chemidoc XRS system (Bio-Rad).

For Streptavidin-HRP blotting, membranes were blocked 10% biotin-free fetal bovine serum (FBS, Fisher Scientific, Cat# 26-400-044) in TBST solution overnight at 4°C with mild agitation. Then membranes were incubated in streptavidin-horseradish peroxidase (1:50000, Life Technologies, Cat# 21130) for one hour at 4°C with mild agitation. After 6 × 15 mins intense wash in TBST solution, the blots were developed and imaged.

Image Lab (Bio-Rad) and Fiji (NIH) were used for image analyses.

### In-gel digestion

After CBB staining, a gel was cut into 11 slices based on protein molecular weight ladder (Biorad, Cat # 1610394) **(Fig. 2H)**. To separate major Shank3 protein isoforms into different gel slices, gel slices A and B were cut based on the observed Shank3 molecular weight from the WB analysis of WT SPN samples **(Fig. 4B)**. Slice A includes top two Shank3 bands. Slice B only contains the third Shank3 band.

We performed in-gel digestions based on a widely-used protocol ^37^. Briefly, each gel slice was put into a microcentrifuge tube and added 500 μl neat ACN to shrink the gel for 10 mins. After removing ACN, DDT solution (10 mM DTT in 100 mM ammonium bicarbonate) was added. The tubes were incubated at 56°C for 30 mins. 500 μl neat ACN was added to remove extra DDT. Then, the gel slices were incubated with IAA (at RT) in dark for 20 mins. After removing extra IAA, the gel slices were saturated with trypsin (13 ng/μl in 10 mM ammonium bicarbonate containing 10% ACN) and incubated at 37°C overnight with intensive agitation. Peptides were extracted from gel slices by incubating with extraction solution (5% formic acid/ACN (1:2 vol/vol)) and vacuum centrifuged to dryness then desalted using ZipTips.

### On-bead digestion

We performed on-beads digestion based on previous reported protocol ^10^. After overnight incubation with striatal homogenates, NeutrAvidin beads were rinsed for five times in one ml lysis buffer (6 M Guanidine, 50 mM HEPES, pH8.5), then added one ml lysis buffer.

Dithiothreitol (DTT, DOT Scientific Inc, Cat # DSD11000) was applied to a final concentration of 5 mM. After incubation at RT for 20 mins, iodoacetamide (IAA, Sigma-Aldrich, Cat # I1149) was added to a final concentration of 15 mM and incubated for 20 mins at RT in the dark. Excess IAA was quenched with DTT for 15 mins. Samples were diluted with buffer (100 mM HEPES, pH 8.5, 1.5 M Guanidine), and digested for three hrs with Lys-C protease (1:100, ThermoFisher Scientific, Cat # 90307_3668048707) at 37°C. Trypsin (1:100, Promega, Cat # V5280) was then added for overnight incubation at 37°C with intensive agitation (1000 rpm). The next day, reaction was quenched by adding 1% trifluoroacetic acid (TFA, Fisher Scientific, O4902-100).

The samples were desalted using HyperSep C18 Cartridges (Thermo Fisher Scientific, Cat # 60108-301) and vacuum centrifuged to dry.

### Tandem Mass Tag (TMT) labeling

Our protocol was based on previously reported methods ^38^. C18 column-desalted peptides were resuspended with 100 mM HEPES pH 8.5 and the concentrations were measured by micro BCA kit (Fisher Scientific, Cat # PI23235). For each sample, 25 μg of peptide labeled with TMT reagent (0.4 mg, dissolved in 40 μl anhydrous acetonitrile, Thermo Fisher Scientific, Cat # 90111) and made at a final concentration of 30% (v/v) acetonitrile (ACN). Following incubation at RT for 2 hrs with agitation, hydroxylamine (to a final concentration of 0.3% (v/v)) was added to quench the reaction for 15 min. For 3-plex TMT experiments, TMT-tagged samples were mixed at a 1:1:1 ratio. For 10-plex TMT experiments, TMT-tagged samples were mixed at a 1:1:1:1:1:1:1:1:1:1 ratio. Combined sample was vacuum centrifuged to dryness, resuspended, and subjected to HyperSep C18 Cartridges.

### Peptide fractionation

We performed the strong cation exchange (SCX) fractionation for all 3-plex TMT-MS experiments. The desalted TMT-labeled sample was fractionated using Hypersep SCX columns (Thermo Fisher Scientific, Cat # 60108-420). Fractions were eluted twice in 300 μl buffer at increasing ammonium acetate concentrations (20, 50, 100, 500, 1000, 2000 mM ammonium acetate). Speed vacuumed to dryness then desalted by ZipTips (Pierce, Cat # 87784) and again dried down for a second time. For all 10-plex TMT-MS experiments, we used a high pH reverse-phase peptide fractionation kit (Thermo Fisher Scientific, Cat # 84868) to get eight fractions (5.0%, 10.0%, 12.5%, 15.0%, 17.5%, 20.0%, 22.5%, 25.0% and 50% of ACN in 0.1% triethylamine solution). The high pH peptide fractions were directly loaded into the autosampler for MS analysis without further desalting.

### Mass spectrometry

Three micrograms of each fraction or sample were auto-sampler loaded with a Thermo EASY nLC 1000 UPLC pump or UltiMate 3000 HPLC pump onto a vented Acclaim Pepmap 100, 75 um x 2 cm, nanoViper trap column coupled to a nanoViper analytical column (Thermo Fisher Scientific, Cat#: 164570, 3 μm, 100 Å, C18, 0.075 mm, 500 mm) with stainless steel emitter tip assembled on the Nanospray Flex Ion Source with a spray voltage of 2000 V. An Orbitrap Fusion (Thermo Fisher Scientific) was used to acquire all the MS spectral data. Buffer A contained 94.785% H2O with 5% ACN and 0.125% FA, and buffer B contained 99.875% ACN with 0.125% FA. For TMT MS experiments, the chromatographic run was for 4 hours in total with the following profile: 0-7% for 7, 10% for 6, 25% for 160, 33% for 40, 50% for 7, 95% for 5 and again 95% for 15 mins receptively. For GelC-MS^2^, the chromatographic run was for 2 hours in total with the following profile: 2–8% for 6, 8–24% for 64, 24–36% for 20, 36–55% for 10, 55–95% for 10, 95% for 10 mins.

We used a multiNotch MS3-based TMT method to analyze all the TMT samples ^38-40^. The scan sequence began with an MS1 spectrum (Orbitrap analysis, resolution 120,000, 400-1400 Th, AGC target 2×10^5^, maximum injection time 200 ms). MS2 analysis, ‘Top speed’ (2 s), Collision-induced dissociation (CID, quadrupole ion trap analysis, AGC 4×10^3^, NCE 35, maximum injection time 150 ms). MS3 analysis, top ten precursors, fragmented by HCD prior to Orbitrap analysis (NCE 55, max AGC 5×10^4^, maximum injection time 250 ms, isolation specificity 0.5 Th, resolution 60,000).

We used CID-MS2 method for GeLC-MS^2^ experiments as previously described ^41^. Briefly, ion transfer tube temp = 300 °C, Easy-IC internal mass calibration, default charge state = 2 and cycle time = 3 s. Detector type set to Orbitrap, with 60K resolution, with wide quad isolation, mass range = normal, scan range = 300-1500 m/z, max injection time = 50 ms, AGC target = 200,000, microscans = 1, S-lens RF level = 60, without source fragmentation, and datatype = positive and centroid. MIPS was set as on, included charge states = 2-6 (reject unassigned). Dynamic exclusion enabled with n = 1 for 30 s and 45 s exclusion duration at 10 ppm for high and low. Precursor selection decision = most intense, top 20, isolation window = 1.6, scan range = auto normal, first mass = 110, collision energy 30%, CID, Detector type = ion trap, OT resolution = 30K, IT scan rate = rapid, max injection time = 75 ms, AGC target = 10,000, Q = 0.25, inject ions for all available parallelizable time.

### MS data analysis and quantification

Protein identification/quantification and analysis were performed with Integrated Proteomics Pipeline - IP2 (Bruker, Madison, WI. http://www.integratedproteomics.com/) using ProLuCID ^42,43^, DTASelect2 ^44,45^, Census and Quantitative Analysis (For TMT MS experiments). Spectrum raw files were extracted into MS1, MS2 and MS3 (For TMT experiments) files using RawConverter (http://fields.scripps.edu/downloads.php). The tandem mass spectra were searched against UniProt mouse protein database (downloaded on 10-26-2020) ^46^ and matched to sequences using the ProLuCID/SEQUEST algorithm (ProLuCID version 3.1) with 5 ppm peptide mass tolerance for precursor ions and 600 ppm for fragment ions. The search space included all fully and half-tryptic peptide candidates within the mass tolerance window with no-miscleavage constraint, assembled, and filtered with DTASelect2 through IP2. To estimate peptide probabilities and false-discovery rates (FDR) accurately, we used a target/decoy database containing the reversed sequences of all the proteins appended to the target database ^47^. Each protein identified was required to have a minimum of one peptide of minimal length of six amino acid residues; however, this peptide had to be an excellent match with an FDR < 1% and at least one excellent peptide match. After the peptide/spectrum matches were filtered, we estimated that the peptide FDRs were ≤ 1% for each sample analysis. Resulting protein lists include subset proteins to allow for consideration of all possible protein forms implicated by at least two given peptides identified from the complex protein mixtures. Then, we used Census and Quantitative Analysis in IP2 for protein quantification of TMT MS. experiments and protein quantification was determined by summing all TMT report ion counts. TMT MS data were normalized using with a build-in method in IP2. For quantification of phosphor-peptides, we calculated the reporter ion intensities from the phosphorylated and unmodified peptides within each protein using a compositional algorithm ^48^ to minimize distortion of the data. Briefly, for every phosphor-protein in each TMT channel, the total TMT reporter ion intensities of all peptides were added up to exactly 2,000,000 (missing values were treated as 0). Renormalized value for each peptide is calculated by the following formula:

TMT_x_-Pi, the reporter ion intensity of peptide i inTMT x channel.

Spyder (MIT, Python 3.7, libraries, ‘pandas’, ‘numpy’, ‘scipy’, ‘statsmodels’ and ‘bioinfokit’) was used for data analyses. RStudio (version, 1.2.1335, packages, ‘tidyverse’, ‘pheatmap’) was used for data virtualization. The Database for Annotation, Visualization and Integrated Discovery (DAVID) (https://david.ncifcrf.gov/) was used for protein functional annotation analysis.

### SPN postsynaptic proteome library

Generation of training dataset: SynaptomeDB_Postsynaptic protein database ^49^ contains a broad range of postsynaptic proteins mostly discovered by proteomic methods. This database has a deep coverage of the postsynaptic proteome but also contains some misclassified presynaptic proteins. SynGO ^50^ is a smaller database and only contains experimentally validated synaptic proteins. Therefore, we combined these two databases to generate a large postsynaptic protein training dataset **(Fig. S3C)**. We first separated presynaptic (SynGO_Presynapse) and postsynaptic proteins (SynGO_Postsynapse) in SynGO. For proteins expressed in both pre- and postsynaptic compartments, we considered them as postsynaptic proteins. SynGO_Presynapse

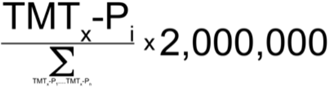

contains proteins which were only expressed in presynaptic compartments. Then we filtered the SynaptomeDB_Postsynaptic protein database with SynGO_Presynapse to remove presynaptic proteins and got SynaptomeDB_Postsynaptic protein_cleaned. We then combined SynaptomeDB_Postsynaptic protein_cleaned and SynGO_Postsynapse to form the final postsynaptic protein training dataset.

Random forest classifier: we used Python library ‘sklearn’ to build our classifier. Proteins were required to have two or more quantified peptides to be considered. For multiple protein isoforms assigned to one gene name, only the isoform with highest total TMT reporter ion intensity was considered. The following features of each protein were used for random forest classifications. They are ‘spec count’, ‘sequence coverage’, ‘molecular weight’, ‘isoelectric point’, ‘log_10_(total peptide intensity/protein length)’, ‘Fold_preBirA* TMT channel/cytoBirA* TMT channel_’, ‘Fold_postBirA* TMT channel/cytoBirA* TMT channel_’, ‘Normalized value in preBirA* TMT channel’, ‘Normalized value in cytoBirA* TMT channel’ and ‘Normalized value in postBirA* TMT channel’. Normalized values were calculated based on a previously reported way ^10^. Briefly, for each TMT channel, the median TMT reporter ion intensity of cytosol and ER proteins was used as the normalization factor. Cytosol and ER proteins were selected based on their GO annotations. Only proteins which were classified as postsynaptic proteins at least twice in two independent 3-plex TMT MS experiments were selected into SPN postsynaptic proteome library. RStudio was used to plot ROC curves.

### Shank3 protein isoform profiling

Standard Shank3 tryptic peptides: for each Shank3 isoform reported in UniProt database (A, B, C1, C3, D1, D2, E1, D2 and F), theoretical Shank3 tryptic peptides were generated by using an online tool, Fragment Ion Calculator (http://db.systemsbiology.net:8080/proteomicsToolkit/FragIonServlet.html). Only fully tryptic Shank3 peptides were included. We removed the peptides whose length is > 35 or < 8 amino acids (AA) because these peptides were not identified in any of our MS experiments.

*Shank3* gene exon mapping: *Shank3* gene exons (NM_021423.4) were linked together in order (from exon 1 to 22). The protein coding sequence was determined by aligning to Shank3A cDNA, which was in-silico (by SnapGene Viewer, 6.0.2) translated into AA sequences. The N-terminal portion of Shank3B, C3, C4, D1, D2, E1, E2 proteins are encoded by alternative spliced transcripts *(25)*. We in silico translated these mRNA transcripts and mapped our peptides to these sequences. For each gel slice, we only considered the Shank3 protein isoforms which match the molecular weight (MW). For example, the theoretical MW of Shank3A is ∼ 185,397 Kda. Therefore, Shank3A should only be detected in gel piece A. Then, we compared Shank3 tryptic peptide sequences with exon-matched and AST-derived AA sequences. In this way, we mapped each Shank3 tryptic peptide into its corresponding exons and ASTs. For peptides that cover AA sequences from two exons A and B (exon-exon junctions), we labeled them as ‘A & B’.

*Shank3* gene exon detection-frequency table: based on exon-mapping results, the detection frequency of each exon (including AST) or exon-joint was calculated by these formulas.

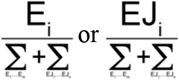

E_i_, number of detection events of exon (or AST) i. EJ_i_, detection times of exon-joint i. In total, m exons (including AST) and n exon-joints were identified. In this way, for each GeLC-MS^2^ experiment, we generated a *Shank3* gene exon detection-frequency table. We also generated standard *Shank3* gene exon detection-frequency table for each Shank3 isoform reported in UniProt database. Then we used tSNE plots to examine the similarity between all *Shank3* gene exon detection-frequency tables. R packages ‘readr’ and ‘Rtsne’ were used to generate tSNE plots.

## Data and codes availability

We will provide the source codes and raw MS data upon acceptance.

## Acknowledgments

Imaging work was performed at the Northwestern University Center for Advanced Microscopy generously supported by NCI CCSG P30 CA060553 awarded to the Robert H Lurie Comprehensive Cancer Center. We would like to thank Farida Vadimovna Korobova, Lennell Reynolds Jr and Wensheng Liu for their help with imaging, Vahri Beaumont for image analysis and Peter Penzes, Jones Griffith Parker and Savas lab members for proofreading this manuscript.

## Funding

Y.Z.W was supported by the Brain & Behavior Research Foundation (2018 Young Investigator Grant, 27793). This research was also supported by an Individual Biomedical Research Award from The Hartwell Foundation to J.N.S. and a grant from CHDI foundation to D.J.S and J.N.S.

## Author contributions

Y.Z.W, D.J.S and J.N.S designed the experiments. Y.Z.W, J.N.S and D.J.S wrote the manuscript.

Y.Z.W and T.P.R performed IHC and EM experiments. Y.Z.W performed all biochemical and MS experiments. T.P.R performed all electrophysiological experiments and related data analysis.

Y.Z.W performed the data analyses for IHC, EM, biochemical and MS experiments.

## Competing interests

The authors declare no competing financial interests.

## Notes

### Competing Interest Statement

The authors have declared no competing interest.

