## Extended data figures for "Neuron-specific protein expression at striatal excitatory synapses"

### Extended data figures and figure legends

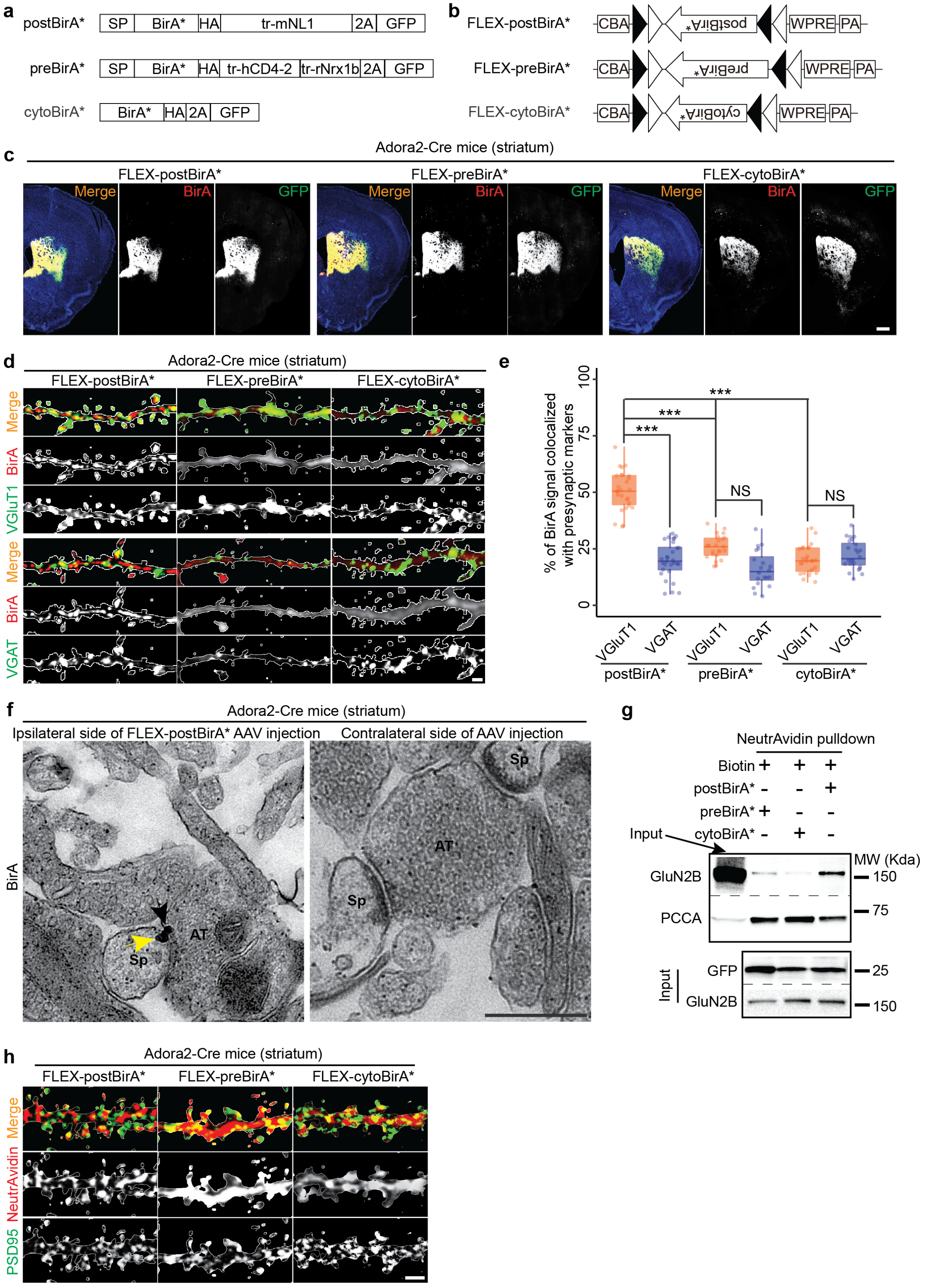

**Extended Data Fig. 1 Confirmation that postBirA* is enriched at postsynaptic sites in striatal iSPNs.**

**(a)** PostBirA* and preBirA* and cytoBirA* control probe constructs. SP, signal peptide, tr-mNL1, truncated mouse Nlgn1, 2A, T2A sequence, tr-hCD4-2, truncated human CD4-2, tr-rNrx1b, truncated rat Nrxn1b.

**(b)** Design of FLEX-postBirA* and control probes. CBA, chicken beta actin promotor, WPRE, Woodchuck hepatitis virus posttranscriptional regulatory element, PA, poly-A sequence.

**(c)** Expressions of BirA* probes in coronal sections showing expression in dorsal striata of Adora2-cre mice. Scale bar, 0.5 mm.

**(d)** PostBirA* colocalizes with the excitatory presynaptic marker VGluT1 to a significantly greater degree compared to the inhibitory presynaptic marker VGAT. Neither preBirA* nor cytoBirA* specifically colocalize with VGluT1 or VGAT. Scale bar, 2 μm.

**(e)** Quantification of **(d)**. n = 4-6 mice, 3-5 brain slices from each mouse. Student's t-test, *** p-value < 0.001. NS, no significant.

**(f)** Immuno-EM micrograph showing two anti-BirA-gold-silver particles, one in the synaptic cleft side (black arrow) and another inside the spine (yellow arrow) from the striatum receiving FLEX-postBirA* AAVs (left). No anti-BirA-gold-silver particles were observed at synapses in the other striatum that did not receive the AAV injection in the same mouse (right). Sp, spine, AT, axonal terminal. Scale bar, 500 nm.

**(g)** Side-by-side comparison of the affinity purified material from all three probes by WB on the same gel. PCCA is the load control for the purified samples.

**(h)** NeutrAvidin-Alex568 fluorescent signals in iSPNs are punctate in postBirA* and partially colocalize with PSD95. The another two BirA* probes have contrasting spatial distributions that are not well-colocalized with PSD95. Scale bar, 2 μm.

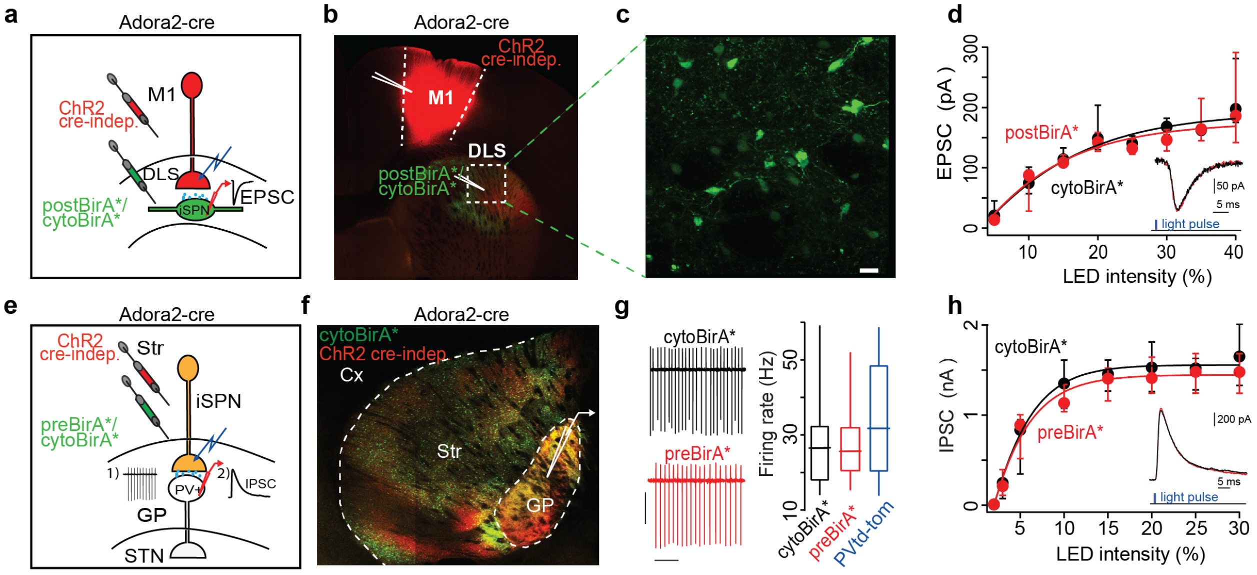


**Extended Data Fig. 2 Unaltered corticostriatal EPSCs and striatopallidal IPSCs in A2a-Cre mice expressing postBirA* or other BirA* probes.**

**(a)** Cartoon showing the experimental design to examine if postBirA* affects corticostriatal EPSCs relative to those expressing cytoBirA*.

**(b)** Low magnification confocal imaging showing Cre-independent channelrhodopsin-2 (ChR2 Cre-indep.) expression in M1 cortex (mCherry) and probes expression in DLS (GFP).

**(c)** High magnification confocal imaging showing co-expression of GFP from FLEX-postBirA* AAVs in iSPNs. Scale bar, 15 µm.

**(d)** Input/output curves for corticostriatal EPSCs at Vm = -80 mV with 15% LED intensity, cytoBirA* = -114 pA (n = 8, 5 mice) vs. postBirA* = -108 pA (n = 9, 6 mice), Bonferroni's multiple comparisons test (p-value > 0.99).

**(e)** Cartoon showing the experimental design to examine if postBirA* affects striatopallidal IPSCs relative to those expressing cytoBirA*.

**(f)** Low magnification confocal imaging showing ChR2 expression in striatum (Str) and globus pallidus (GP) (mCherry) and cytoBirA* expression (GFP).

**(g)** Left, Striatopallidal IPSCs were recorded from presumably parvalbumin positive (PV+) GP neurons at 15-60 Hz. Right, Summary of firing rates (p-value = 0.57, Kruskal-Wallis test), we used historical controls from PV+ neurons recorded in PV tdTomato mice.

**(h)** Input/output curves of striatopallidal IPSCs at Vm = - 40 mV with 15% LED intensity, ctrl2BirA*: 1455 pA (n = 7,4 mice) vs. preBirA*: 1404 pA (n = 8, 5 mice), Bonferroni's multiple comparisons test, p-value > 0.99). Cx, Cortex, Globus pallidus, GP, Str, striatum, DLS, dorsolateral striatum.

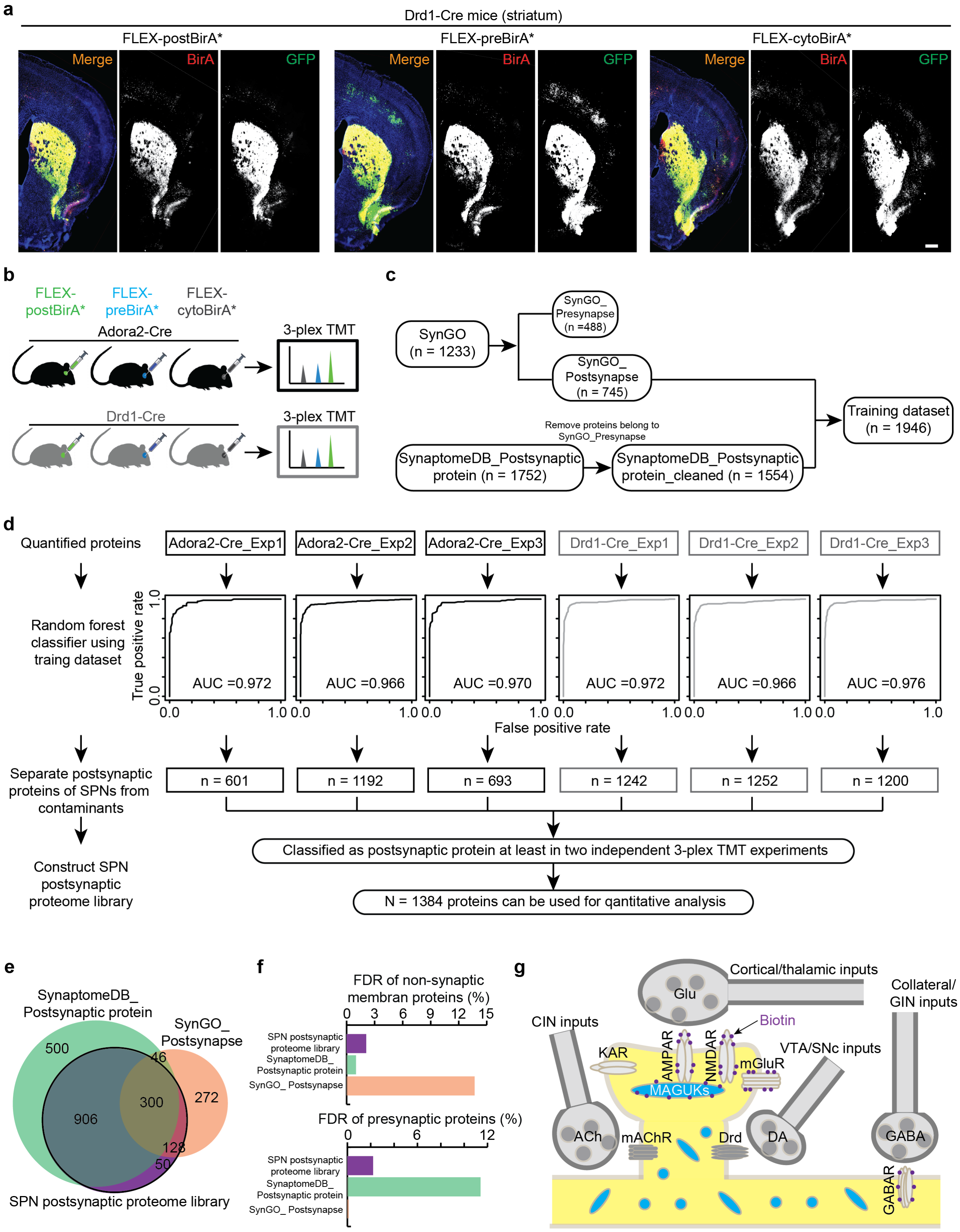
a
Drd1-Cre mice (striatum)
FLEX-postBirA*
FLEX-preBirA*
FLEX-cytoBirA*
Merge
BirA
GFP
b
FLEX-postBirA*
FLEX-preBirA*
FLEX-cytoBirA*
Adora2-Cre
3-plex TMT
Drd1-Cre
3-plex TMT
c
SynGO (n = 1233)
SynGO_ Presynapse (n =488)
SynGO_ Postsynapse (n = 745)
Remove proteins belong to SynGO_Presynapse
SynaptomeDB_Postsynaptic protein (n = 1752)
SynaptomeDB_Postsynaptic protein_cleaned (n = 1554)
Training dataset (n = 1946)
d
Quantified proteins
Adora2-Cre_Exp1
Adora2-Cre_Exp2
Adora2-Cre_Exp3
Drd1-Cre_Exp1
Drd1-Cre_Exp2
Drd1-Cre_Exp3
Random forest classifier using traing dataset
True positive rate
AUC =0.972
AUC =0.966
AUC =0.970
AUC =0.972
AUC =0.966
AUC =0.976
False positive rate
Separate postsynaptic proteins of SPNs from contaminants
n = 601
n = 1192
n = 693
n = 1242
n = 1252
n = 1200
Classified as postsynaptic protein at least in two independent 3-plex TMT experiments
Construct SPN postsynaptic proteome library
N = 1384 proteins can be used for qantitative analysis
e
SynaptomeDB_ Postsynaptic protein
SynGO_ Postsynapse
500
46
300
272
906
128
50
SPN postsynaptic proteome library
f
FDR of non-synaptic membran proteins (%)
SPN postsynaptic proteome library
SynaptomeDB_ Postsynaptic protein
SynGO_ Postsynapse
FDR of presynaptic proteins (%)
g
Cortical/thalamic inputs
Glu
Collateral/ GIN inputs
CIN inputs
Biotin
KAR
AMPAR
NMDAR
mGluR
VTA/SNc inputs
MAGUKs
ACh
mAChR
Drd
DA
GABA
GABAR

**Extended Data Fig. 3 Determine SPN postsynaptic proteome library.**

**(a)** Representative BirA* AAV GFP and BirA expression in dorsal striata of Drd1-cre mice. Scale bar, 0.5 mm.

**(b)** Experimental design for determining high-confidence postsynaptic compartment proteins biotin-tagged by postBirA* in SPNs. Three independent 3-plex TMT-MS experiments, each channel was pooled from three mice for each Cre line. N = 54 total mice.

**(c)** Generation of training dataset by combination of postsynaptic proteins reported in SynaptomeDB and SynGO databases.

**(d)** Full set of bioinformatic analyses to determine which proteins can be used for quantitative analysis in further experiments. ROC curves of six 3-plex TMT-MS datasets.

**(e)** Coverage of SPN postsynaptic proteome library.

**(f)** For SPN postsynaptic proteome library, the false-discovery rate (FDR) of non-synaptic membrane proteins is 2.1%. The FDR of presynaptic proteins is 2.2%.

**(g)** Cartoon schematic illustrating selected postsynaptic proteins biotinylated by postBirA* in SPNs. Glu, glutamate, Ach, acetylcholine, DA, dopamine, GIN, GABAergic interneuron, CIN, cholinergic interneuron, VTA, ventral tegmental area, SNc, substantia nigra pars compacta.

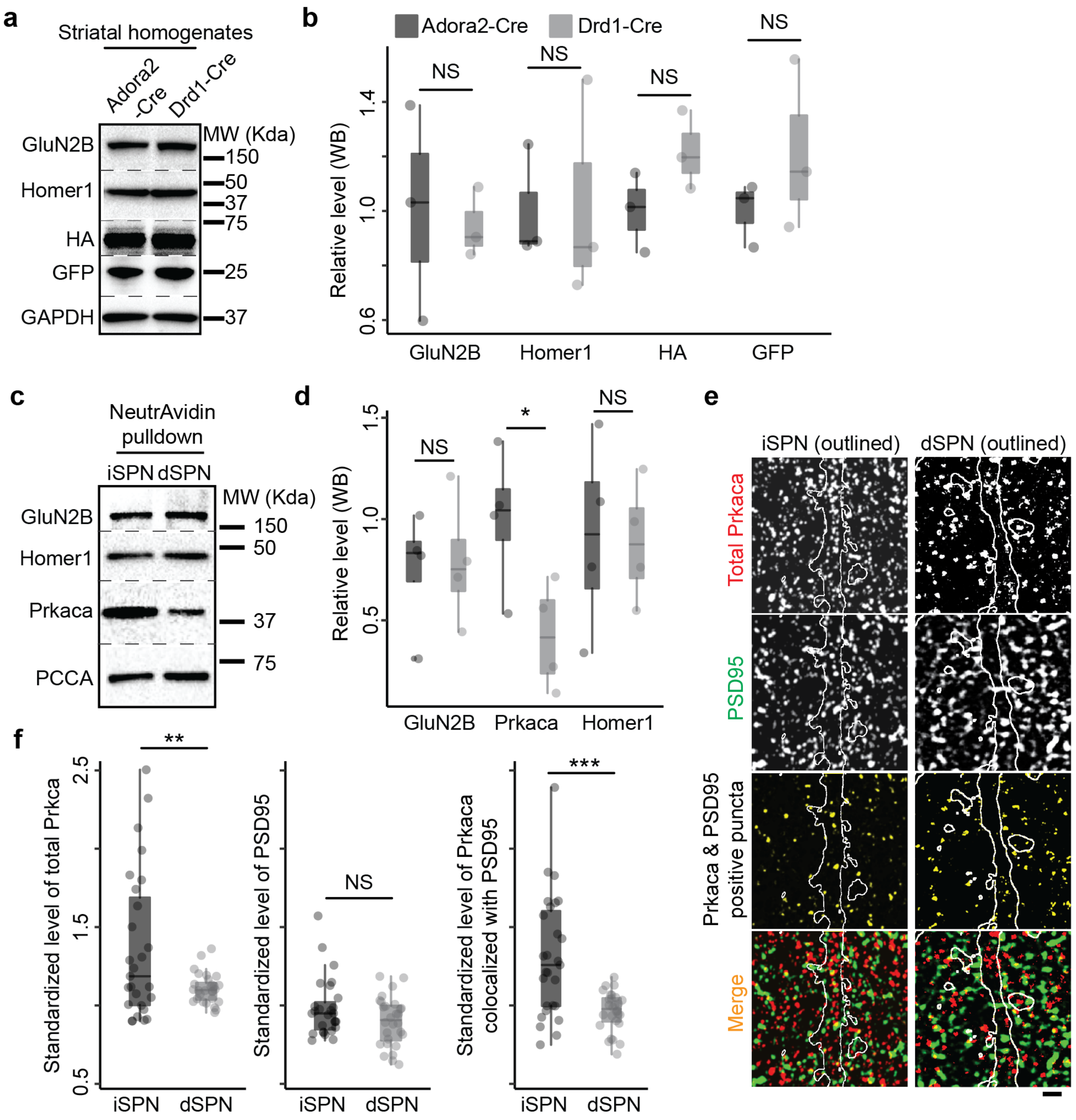


**Extended Data Fig. 4 Validation of TMT-MS data.**
**(a)** The expression levels of postBirA* were similar in Adora2- and Drd1- Cre striata, based on HA and GFP WB.
**(b)** Quantification of **(a)**, n = 3 mice.
**(c)** Prkaca level is significantly higher in affinity purified material from Adora2-Cre striata expressing postBirA* compared to Drd1-Cre.
**(d)** Quantification of **(c)**, protein levels are normalized to PCCA. n = 4 mice per genotype.
**(e)** PSD95-Prkaca colocalized signals are significantly more abundant in iSPN compared to dSPN dendrites. White outline indicates GFP signal. Scale bar, 1 μm.
**(f)** Quantification of **(e)**, n = 5 mice, 4-6 brain slices from each mouse.

**(b, d & f)** Student's t-test, * p-value < 0.05, ** p-value < 0.01, *** p-value < 0.001, NS, not significant.

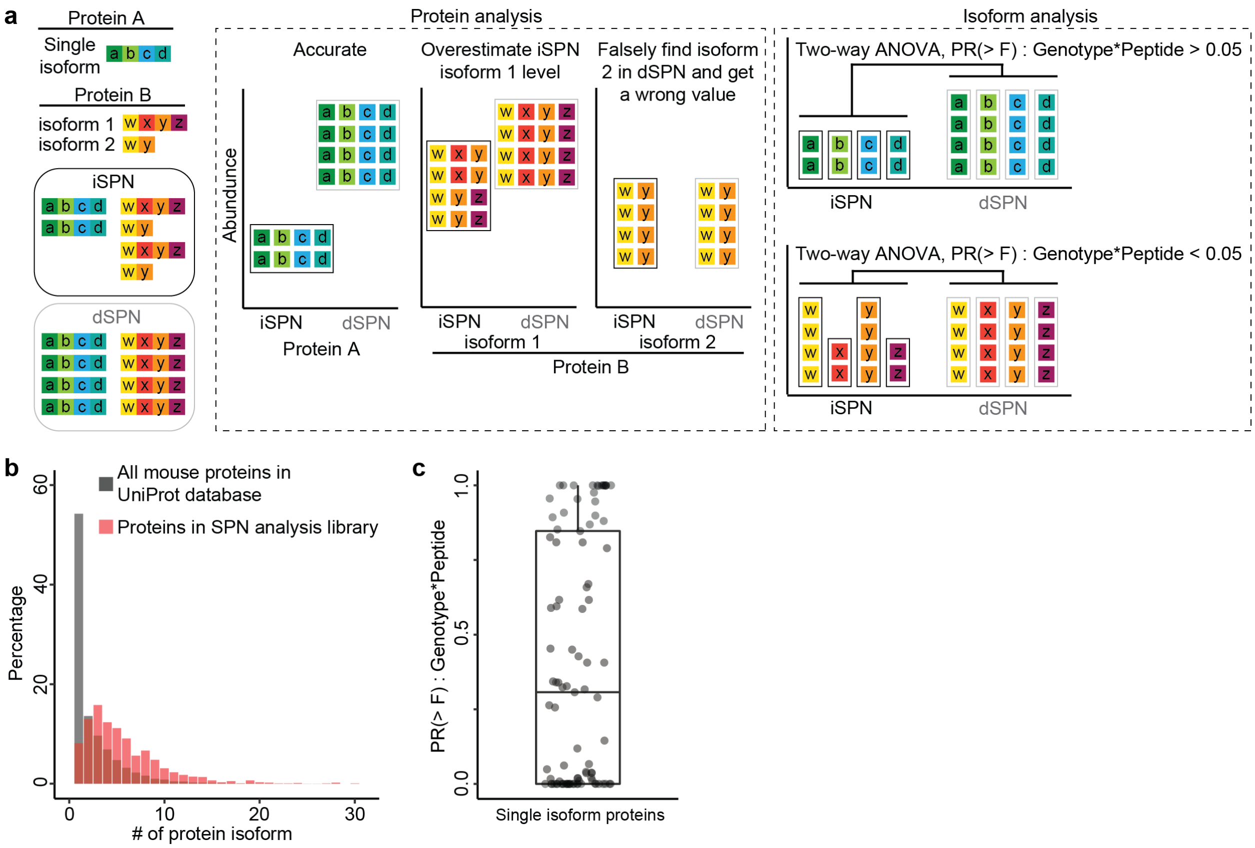


**Extended Data Fig. 5 Logic map of isoform analysis strategy of iSPN and dSPN proteomes by two-way ANOVA.**

**(a)** Comparison of protein analysis and isoform analysis. In protein analysis, the abundance of a protein is calculated based on the relative abundance of all the quantified peptides. This quantification method works very well when the quantified protein only has one isoform (like protein A) or if you consider protein groups. However, it is not well suited to quantify multi-isoform proteins (e.g., protein B). Two-way ANOVA (factors: peptide, genotype) can tell whether the heteroscedastic variances of all quantified peptides are significantly (PR > F): Genotype * peptide < 0.05) related to genotypes. Therefore, isoform analysis can be used to detect divergent expression at the protein isoform level in SPNs. However, isoform analysis cannot be used for quantifications.

**(b)** Based on UniProt mouse protein database, more than 50% of genes can produce multiple protein isoforms. Interestingly, nearly 90% of genes belonging to our SPN analysis library can produce multiple protein isoforms.

**(c)** The median of (PR > F): Genotype * peptide values of single isoform proteins is 0.307. The mean is 0.392. The data suggest that the genotype didn't contribute to the variances of quantified peptides of single isoform proteins.

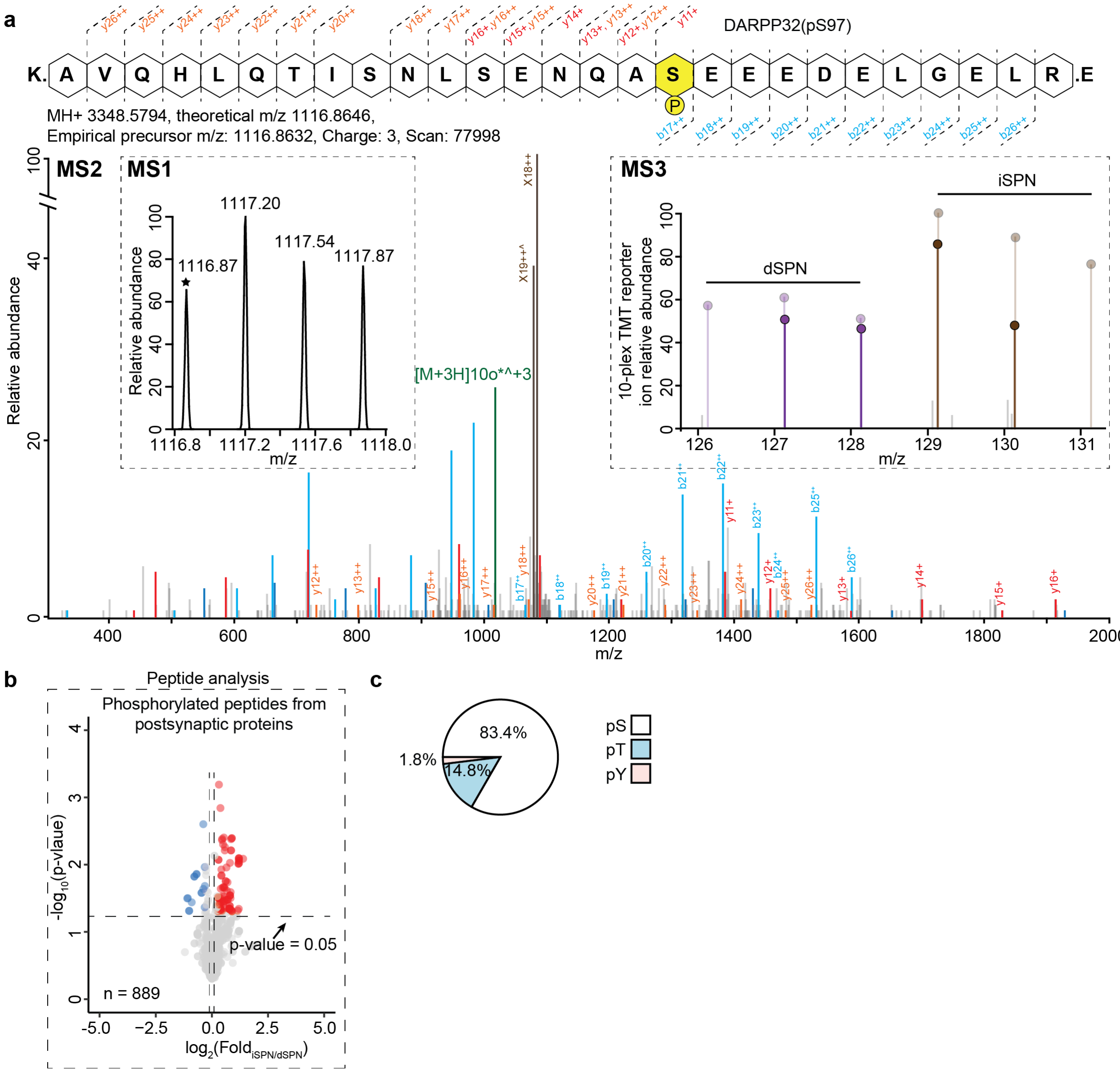


**Extended Data Fig. 6 Differential phosphorylation status of iSPN and dSPN postsynaptic proteins.**

**(A)** MS spectra of the indicated DARPP32 phosphorylated peptide (pS97) acquired from postBirA* TMT-MS experiment in WT dSPNs and iSPNs. MS1 peak selected for MS2 is indicated with a star. For MS2 spectra assigned b (dark blue +, light blue ++) and y (red +, orange ++) fragment ions are indicated and those containing phosphorylated Serine 97 are labeled. MS3 spectra shows 10-plex TMT reporter ion intensities from dSPNs (purple) and iSPNs (brown).

**(B)** Peptide analysis showing that the overall level of postsynaptic protein phosphorylation in iSPNs is significantly higher than in dSPNs at the phopshopeptide level. Fisher's combined p-value = 1.84E-89.

**(C)** Pie charts showed the amino acid compositions of total set of identified phosphorylation sites.

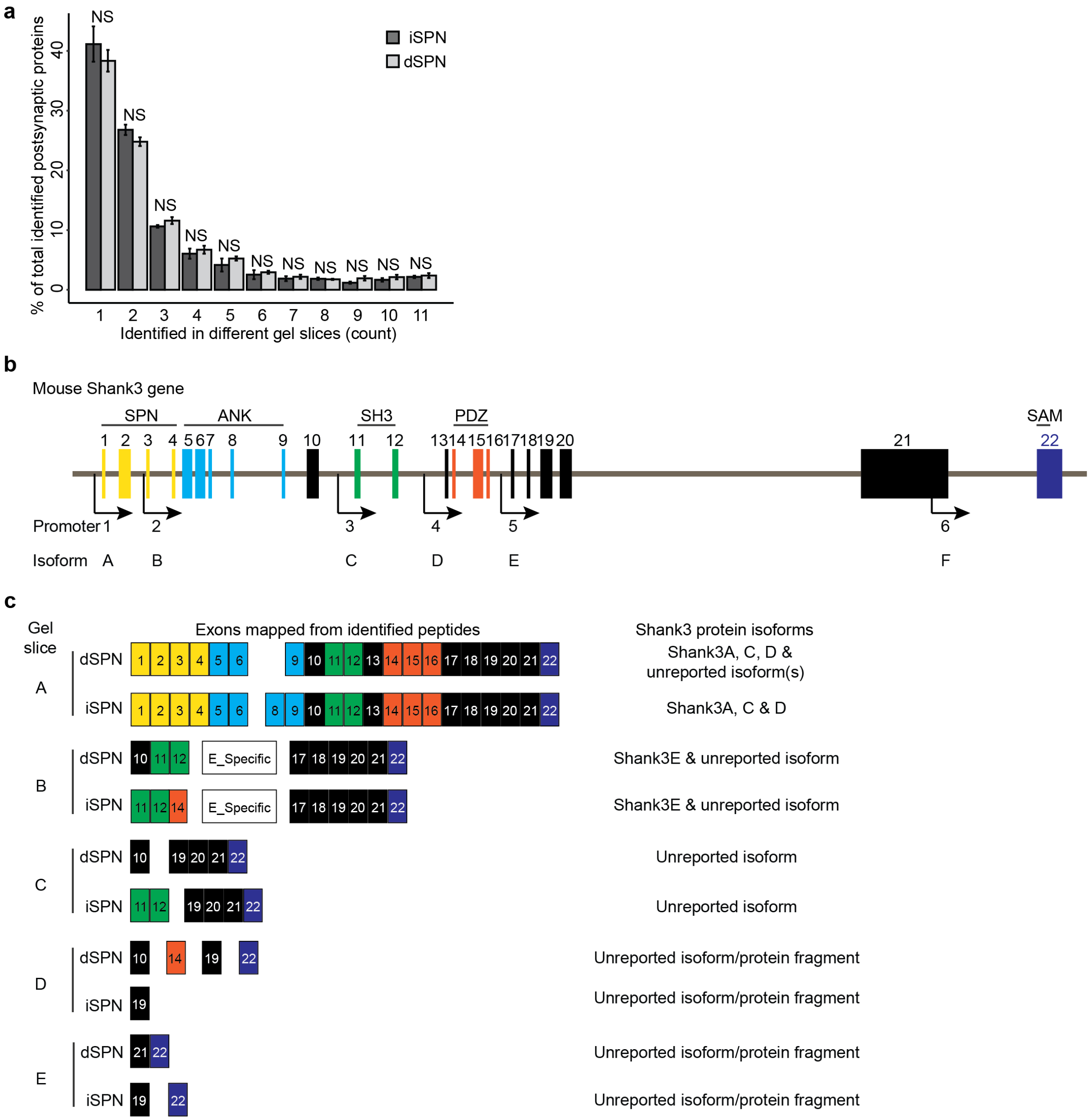


**Extended Data Fig. 7 All Shank3 peptides identified in GeLC-MS[2] experiments mapped to *Shank3* gene exons.**

**(a)** Summary of GelC-MS2 analysis. Nearly 60% of identified proteins are detected in multiple gel pieces. N = 3 mice for each genotype.

**(b)** Mouse Shank3 gene (Gene ID: 58234) has 22 exons and six known promotors. Alternative usage of promotors directs the expressions of six major Shank3 protein isoforms. ANK, ankyrin repeats domain, SH3, src 3 domain, PRO, proline-rich domain, SAM, sterile α motif domain.

**(c)** Peptides identified in each gel pieces were mapped to *Shank3* gene exons (NM_021423.4).

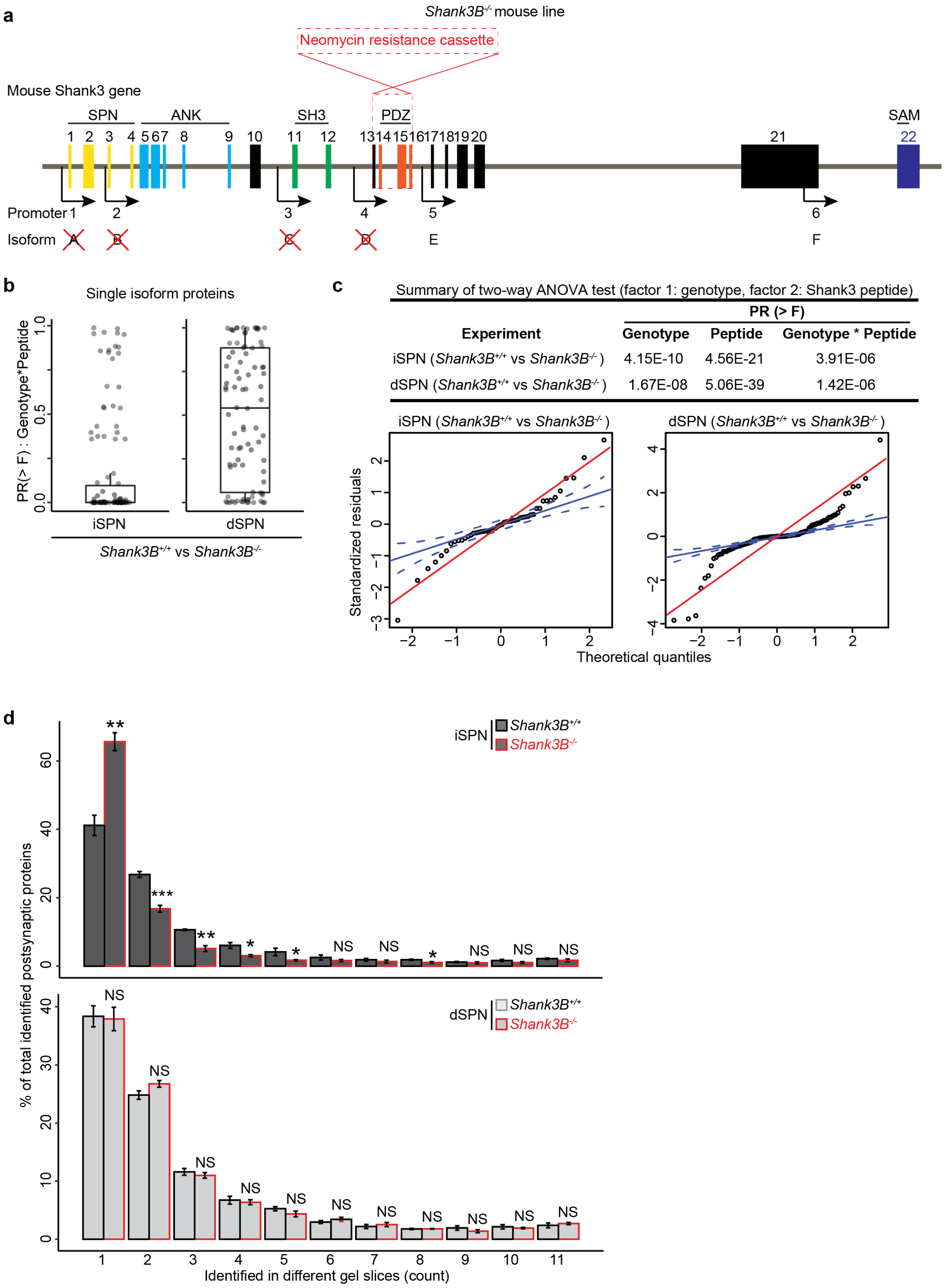


Summary of two-way ANOVA test (factor 1: genotype, factor 2: Shank3 peptide)

| Experiment | PR (> F) Genotype | Peptide | Genotype * Peptide |
|---|---|---|---|
| iSPN (*Shank3B*+/+ vs *Shank3B*-/-) | 4.15E-10 | 4.56E-21 | 3.91E-06 |
| dSPN (*Shank3B*+/+ vs *Shank3B*-/-) | 1.67E-08 | 5.06E-39 | 1.42E-06 |

**Extended Data Fig. 8 *Shank3* gene exons 13-16 deletion divergently altered Shank3 protein isoforms in iSPN and dSPN postsynaptic compartments.**

**(a)** In *Shank3B*$^{-/-}$ mouse line, exons 13-16 were replaced with neomycin resistant cassette. It disrupted the expressions of Shank3A, B, C & D.

**(b)** The (PR > F): Genotype * peptide values of single isoform proteins two 10-plex TMT experiments. For *Shank3B*$^{+/+}$ vs *Shank3B*$^{-/-}$ iSPNs experiment, the median is 0.0007, the mean is 0.142. For *Shank3B*$^{+/+}$ vs *Shank3B*$^{-/-}$ dSPNs, the median is 0.541, the mean is 0.504.

**(c)** Q-Q plots showing that the variances of Shank3 peptides in *Shank3B*$^{-/-}$ iSPNs and dSPNs are heteroscedastic, which suggests that Shank3 protein isoforms are dissimilar in these two neuron types.

**(d)** Summary of GelC-MS2 experiments. Notably, *Shank3* gene exons 13-16 deletion significantly increased the percentage of proteins only detected in single gel piece, which suggests that protein-isoform complexity was reduced in *Shank3B*$^{-/-}$ iSPN postsynaptic compartments. N = 3 mice per genotype.
