## Supplementary information for "Neuron-specific protein expression at striatal excitatory synapses"

**This file included:**

Legends for Tables S1 to S5

**Table S1. SPN postsynaptic proteome library**

Full list of proteins included in the SPN postsynaptic proteome library.

**Table S2. Complete list of quantified postsynaptic proteins in iSPNs vs dSPNs postBirA* TMT-MS analyses**

Sheet1: Summary of protein analysis. The indicated intensity values for each biological replicate are calculated from the normalized average of the TMT reporter ion intensities from all of the corresponding peptides spectral matches for each protein. BR = biological replicate (individual mice). Student’s t-test. SigDn, significantly down-regulated in iSPN postsynaptic compartments (p-value < 0.05, fold_dSPN/iSPN_ > 1.2). SigUp, significantly up-regulated in iSPN postsynaptic compartments (p-value < 0.05, fold_iSPN/dSPN_ > 1.2). NonSig, no significant difference.

Sheet2: Summary of isoform analysis. Two-way ANOVA test. Similar, (PR > F): Genotype * peptide value > 0.05, dissimilar, (PR > F): Genotype * peptide value < 0.05.

Sheet3: Summary of quantified phosphorylated peptides. The values indicated represent reporter ion intensities for each biological replicate, calculated from the normalized average of the TMT reporter ion intensities from all of the corresponding peptides spectral matches for each phospho-peptides. Student’s t-test.

**Table S3. List of all Shank3 peptides in GeLC-MS^2^ experiments from WT iSPNs and dSPNs.**

Sheet 1: All Shank3 peptides identified in gel piece A.

Sheet 2: All Shank3 peptides identified in gel piece B.

Sheet 3: All Shank3 peptides identified in gel piece C.

Sheet 4: All Shank3 peptides identified in gel piece D.

Sheet 5: All Shank3 peptides identified in gel piece E.

Shank3 peptide sequence, mapped to *Shank3* gene exons, XCorr, ΔCN scores, confidence (Conf%), parts per million (PPM), spectral count (SpecCount) and retention time (RetTime) from Sequest / ProLucid are indicated.

**Table S4. Complete list of quantified postsynaptic proteins in *Shank3B^-/-^* iSPNs and dSPNs compared to WT controls from postBirA* TMT-MS analyses**

Sheet 1: Summary of all quantified postsynaptic proteins from postBirA* TMT-MS analysis of iSPNs in *Shank3B^-/-^* compared to WT control mice. Protein analysis. Student’s t-test. SigDn, significantly down-regulated in *Shank3B^-/-^* iSPN postsynaptic compartments (p-value < 0.05, fold_Shank3B+/+ /Shank3B-/-_ > 1.2). SigUp, significantly up-regulated in *Shank3B^-/-^* iSPN postsynaptic compartments (p-value < 0.05, fold_Shank3B-/-/Shank3B+/+_ > 1.2). NonSig, no significant difference.

Sheet 2: Summary of all quantified postsynaptic proteins from postBirA* TMT-MS analysis of dSPNs in *Shank3B^-/-^* compared to WT control mice. Protein analysis. Student’s t-test. SigDn, significantly down-regulated in *Shank3B^-/-^* dSPN postsynaptic compartments (p-value < 0.05, fold_Shank3B+/+ /Shank3B-/-_ > 1.2). SigUp, significantly up-regulated in *Shank3B^-/-^* dSPN postsynaptic compartments (p-value < 0.05, fold_Shank3B-/-/Shank3B+/+_ > 1.2). NonSig, no significant difference.

Sheet 3: Summary of isoform analysis on *Shank3B^+/+^* vs *Shank3B^-/-^* iSPN TMT-MS experiment.

Two-way ANOVA test. Similar, (PR > F): Genotype * peptide value > 0.05, dissimilar, (PR > F): Genotype * peptide value < 0.05.

Sheet 4: Summary of isoform analysis on *Shank3B^+/+^* vs *Shank3B^-/-^* dSPN TMT-MS experiment. Two-way ANOVA test. Similar, (PR > F): Genotype * peptide value > 0.05, dissimilar, (PR > F): Genotype * peptide value < 0.05.

**Table S5. List of all Shank3 peptides in GeLC-MS^2^ experiments from *Shank3B^-/-^*** **iSPNs and dSPNs.**

Sheet 1: All Shank3 peptides identified in gel piece A.

Sheet 2: All Shank3 peptides identified in gel piece B.

Sheet 3: All Shank3 peptides identified in gel piece C.

Sheet 4: All Shank3 peptides identified in gel piece F.

Shank3 peptide sequence, mapped to *Shank3* gene exons, XCorr, ΔCN scores, confidence (Conf%), parts per million (PPM), spectral count (SpecCount) and retention time (RetTime) from Sequest / ProLucid are indicated.
